# Impact of histone N-terminal domains and linker DNA on H2A.Z deposition by yeast SWR1C

**DOI:** 10.64898/2026.05.01.722189

**Authors:** Yonca B. Karadeniz, Craig L. Peterson

**Affiliations:** Program in Molecular Medicine, University of Massachusetts Chan Medical School, Worcester, MA USA; Interdisciplinary Graduate Program, University of Massachusetts Chan Medical School

**Keywords:** Chromatin remodeling, SWR1C, histone acetylation, FRET, NuA4, dinucleosomes, linker DNA

## Abstract

ATP-dependent chromatin remodeling enzymes play a central role in governing the essential functions of the genome, and deficiencies in their catalytic function often lead to developmental disorders or disease. While most remodelers use the energy of ATP hydrolysis to either mobilize nucleosomes or evict histone octamers, yeast SWR1C catalyzes the ATP-dependent replacement of nucleosomal H2A/H2B dimers with variant H2A.Z/H2B dimers. The H2A.Z histone variant is conserved in all eukaryotes where it plays key roles in regulating gene transcription, DNA repair, and heterochromatin function. Consequently, the regulation of H2A.Z deposition by SWR1C is central to numerous genomic functions. Here, we use both quantitative fluorescence-based assays and gel-based, electrophoretic methods to dissect the roles of histone tails, histone acetylation, and linker DNA in SWR1C-mediated H2A.Z deposition in vitro. Unlike many other remodeling enzymes, we find that histone tails are largely dispensable for the catalytic activity of SWR1C, and histone acetylation by the NuA4 acetyltransferase has only a minor stimulatory impact on SWR1C activity. In contrast, we confirm previous studies that nucleosome-free, linker DNA stimulates SWR1C activity even under saturating enzyme levels. These insights provide a clearer understanding of the structural and regulatory determinants that guide H2A.Z deposition by SWR1C, offering potential new avenues to investigate its role in chromatin dynamics and genome stability.

## Introduction

Eukaryotic genomes are organized into complex chromatin structures that at the most basic level consist of long, linear arrays of nucleosomes that each contain ∼147bp of DNA wrapped nearly twice around an octamer of the core histones, H2A, H2B, H3, and H4. Canonical histones and their variants are characterized by well-structured histone fold domains and long, unstructured N-terminal domains (also known as histone tails), which provide crucial sites for post-translational modifications (PTMs), such as lysine acetylation. These histone tails protrude from the nucleosome and interact with DNA and protein factors. Chromatin architecture is dynamic in order to meet diverse cellular needs, ranging from the regulation of gene expression, cell cycle regulation, environmental adaptation, development, and to aging (Luger et al., 1999). A key aspect of this regulation is mediated by ATP-dependent chromatin remodelers, which modulate nucleosome positioning and histone composition, playing a critical role in DNA accessibility, transcription, DNA replication, repair, and other genomic processes (Clapier and Cairns, 2009; Clapier et al., 2017).

In the budding yeast, *Saccharomyces cerevisiae*, chromatin is organized into nucleosomal arrays where each nucleosome is separated by ∼18bp of linker DNA. Upstream of mRNA genes, there is typically an additional nucleosome-free region (NFR) of ∼150bp that is flanked by positioned -1 and +1 nucleosomes (Guillemette et al., 2005; Jiang and Pugh, 2009; Kaplan et al., 2009; Abril-Garrido et al., 2023). The NFR contains gene regulatory sequences, and the +1 nucleosome is often overlapping or adjacent to the transcription start site. Notably, nucleosomes that flank the NFR are often enriched for the histone variant H2A.Z, which facilitates inducible gene transcription and plays a global transcriptional role in cells that lack the RNA exosome (Guillemette et al., 2005; Raisner et al., 2005; Jin et al., 2009; Bryll and Peterson, 2022). Understanding the regulation of H2A.Z deposition and how it affects chromatin structure and function are crucial for insights into gene regulation and genome stability.

SWR1C is a multisubunit, megadalton ATP-dependent chromatin remodeling complex that is responsible for the deposition of H2A.Z into yeast chromatin (Krogan et al., 2003; Mizuguchi et al., 2004). SWR1C is a member of the INO80 subfamily of ATP-dependent chromatin remodeling enzymes, but unlike all other remodelers, SWR1C is unable to mobilize nucleosomes along DNA, but rather its activity is limited to the ATP-dependent exchange of nucleosomal H2A/H2B dimers for variant H2A.Z/H2B dimers. The ATPase subunit of SWR1C, Swr1, engages nucleosomal DNA ∼2 helical turns from the nucleosomal dyad, a position called SHL 2. This binding site places the ATPase lobes in close apposition to the protruding histone H4 N-terminal domain, and members of the ISWI, SWI/SNF, and CHD remodeler families are regulated by interactions with this adjacent H4 tail (Yan et al., 2016; Farnung et al., 2017; Liu et al., 2017). Currently, it is unclear if the H4 tail impacts H2A.Z deposition by SWR1C, though one previous study suggested that histone tail domains and their acetylation may promote either SWR1C nucleosome binding or activity (Altaf et al., 2010). Understanding the regulatory mechanisms that control SWR1C is crucial, as chromatin remodelers share conserved functional and regulatory cues across species, providing insights that could lead to therapeutic interventions for chromatin remodeler-related disorders. Despite substantial progress, how SWR1C integrates nucleosomal regulatory cues to tune H2A.Z exchange remains incompletely understood. In particular, the extent to which SWR1C activity depends on histone tail domains and tail acetylation versus extranucleosomal DNA features such as linker DNA— and how these determinants behave in the context of adjacent nucleosomes—has not been systematically defined in a quantitative biochemical framework. Because H2A.Z is preferentially deposited at nucleosomes flanking promoter nucleosome-free regions, defining how linker DNA and nucleosome context modulate SWR1C is central to understanding its targeting and regulation.

In this work, we investigated the ability of SWR1C to deposit H2A.Z in response to different regulatory elements of the nucleosome. We examined how SWR1C activity was impacted by (1) histone tail regions, (2) acetylated histones H2A and H4, (3) linker DNA length, and (4) the role of these chromatin elements in the context of dimeric nucleosome arrays. Surprisingly, we find that histone tails have little impact on SWR1C activity, although SWR1C does require nucleosome-free linker DNA for optimal activity. The requirement for linker DNA is consistent with its preference *in vivo* for depositing H2A.Z adjacent to an NFR. Insights from this research could provide a deeper understanding of chromatin dynamics and inform strategies for targeting chromatin remodeler-related disorders.

## Results

### Histone tails are dispensable for SWR1C-mediated H2A.Z exchange

SWR1C catalyzes the stepwise removal of canonical H2A/H2B dimers from nucleosomes and the subsequent deposition of H2A.Z/H2B dimers (Figure 1A). To investigate whether histone tails are necessary for SWR1C-mediated H2A.Z dimer exchange, we performed a fluorescence-based dimer exchange assay, leveraging Förster resonance energy transfer (FRET) to monitor nucleosome composition over time. The nucleosomal substrates contain a Cy3 fluorophore conjugated to DNA at one edge of the nucleosome, and a Cy5 fluorophore attached to an engineered cysteine residue (H2A-119) within the histone H2A C-terminal domain. Nucleosomes were reconstituted with recombinant histones that contained or lacked the histone N-terminal tail domains (recTailless) or only the N-terminal tail domain of histone H4 (H4 tailless), using 147 bp Widom 601 sequence and 77 bp extranucleosomal linker DNA on one side. The Cy3 and Cy5 fluorophores are within an appropriate distance to function as a FRET pair, such that excitation of the Cy3 donor leads to efficient energy transfer to the Cy5 acceptor, as evidenced by the fluorescence emission peak at 670 nm. The dimer exchange activity of SWR1C is monitored by following the decrease in the 670 nm FRET signal due to eviction of the Cy5-labeled H2A/H2B dimer (Figure 1A).

**Figure 1.**
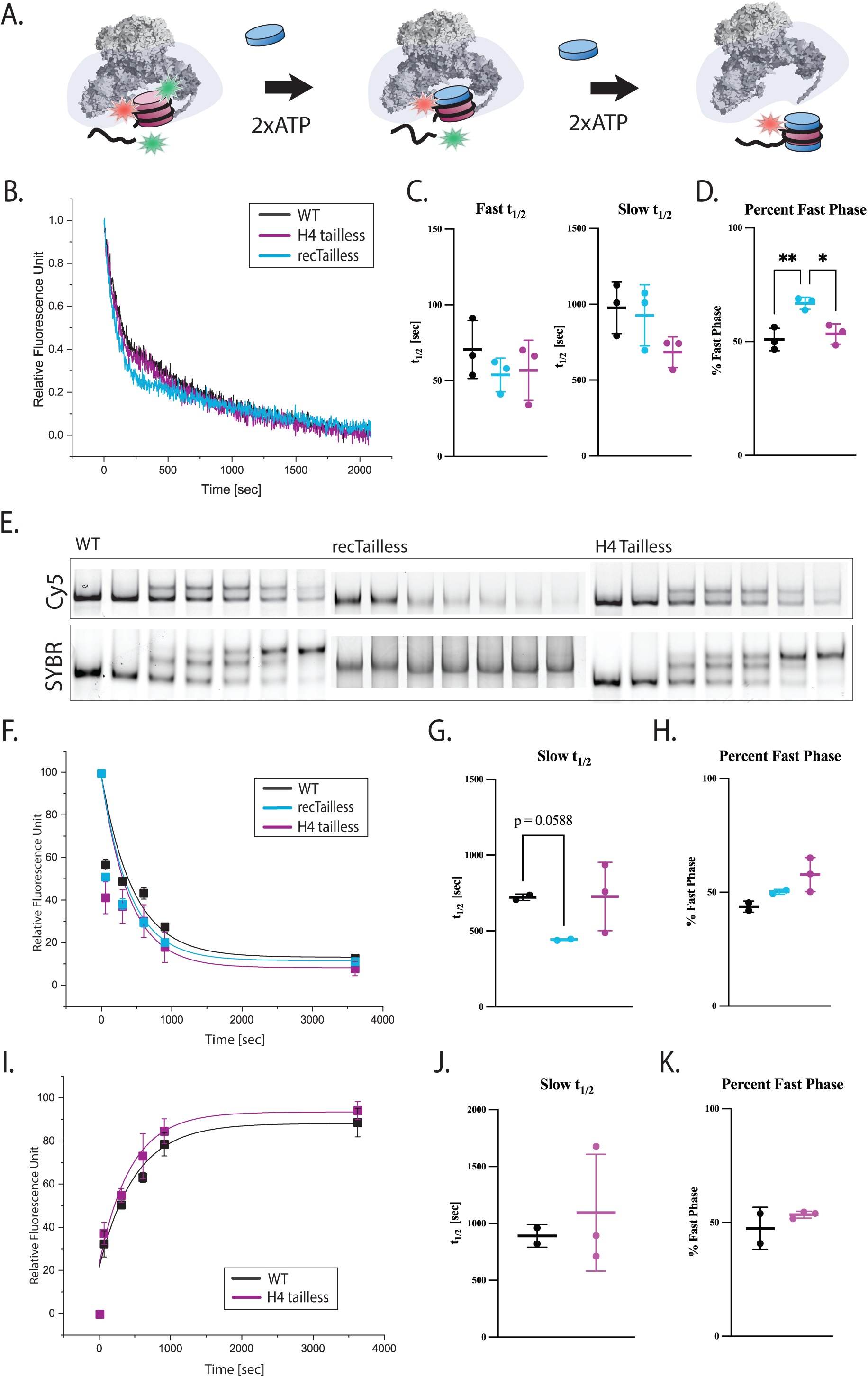
Histone tails are dispensable for H2A.Z deposition. **A.** Schematic of the ensemble FRET-based dimer-exchange (eviction) assay. Nucleosomes contained Cy5-labeled H2A and Cy3-labeled DNA. During SWR1C-catalyzed replacement of Cy5–H2A/H2B dimers, the FRET signal decreases. **B.** Eviction kinetics comparing WT (black) and recombinant tailless (blue) or H4-tailless (red) nucleosomes (*N* = 3). **C–D.** Kinetic parameters from fits to traces in (B): fast and slow half-times (t₁/₂, s) (**C**), and fast-phase fraction (%) (Fast phase amplitude divided by total of fast and slow phase amplitudes) (**D**). **E.** Representative native gel-based dimer-exchange assay under single-turnover conditions. Samples were collected as nucleosome only, time 0, and at 1, 5, 10, 15, and 60 min after ATP addition. SYBR Gold staining (top) reports DNA-containing species; Cy5 fluorescence (bottom) reports Cy5–H2A/H2B–containing species. **F–K.** Quantification of gel-based assays for eviction (**F–H**) and deposition (**I–K**) as indicated (each condition *N* = 2–3). For eviction, ACy5/ACy5 and ACy5/Z species were pooled; for deposition, ACy5/Z and Z/Z species were pooled. Data were scaled from basal to plateau (0–1) for visualization and plotted versus time (s). Kinetic models (one-phase or two-phase association/decay, as indicated) were fit to the unscaled data to obtain t₁/₂ and fast-phase fraction (reported as mean ± SD). Statistics: Welch’s ANOVA with Dunnett’s T3 multiple comparisons was used for panels **C, D, G, H**; Welch’s *t* test was used for **J, K**. For percent fast phase (WT, RecTailless, H4Tailless), Welch’s ANOVA: W(DFn, DFd)=15.16 (2, 3.686), p=0.0167. Dunnett’s T3 pairwise comparisons: WT vs RecTailless t=4.911, df=3.068, p=0.0380; RecTailless vs H4Tailless t=4.510, df=3.265, p=0.0477. All additional comparisons/statistics are reported in the statistics table (Supplemental Table 1).

**Figure 2.**
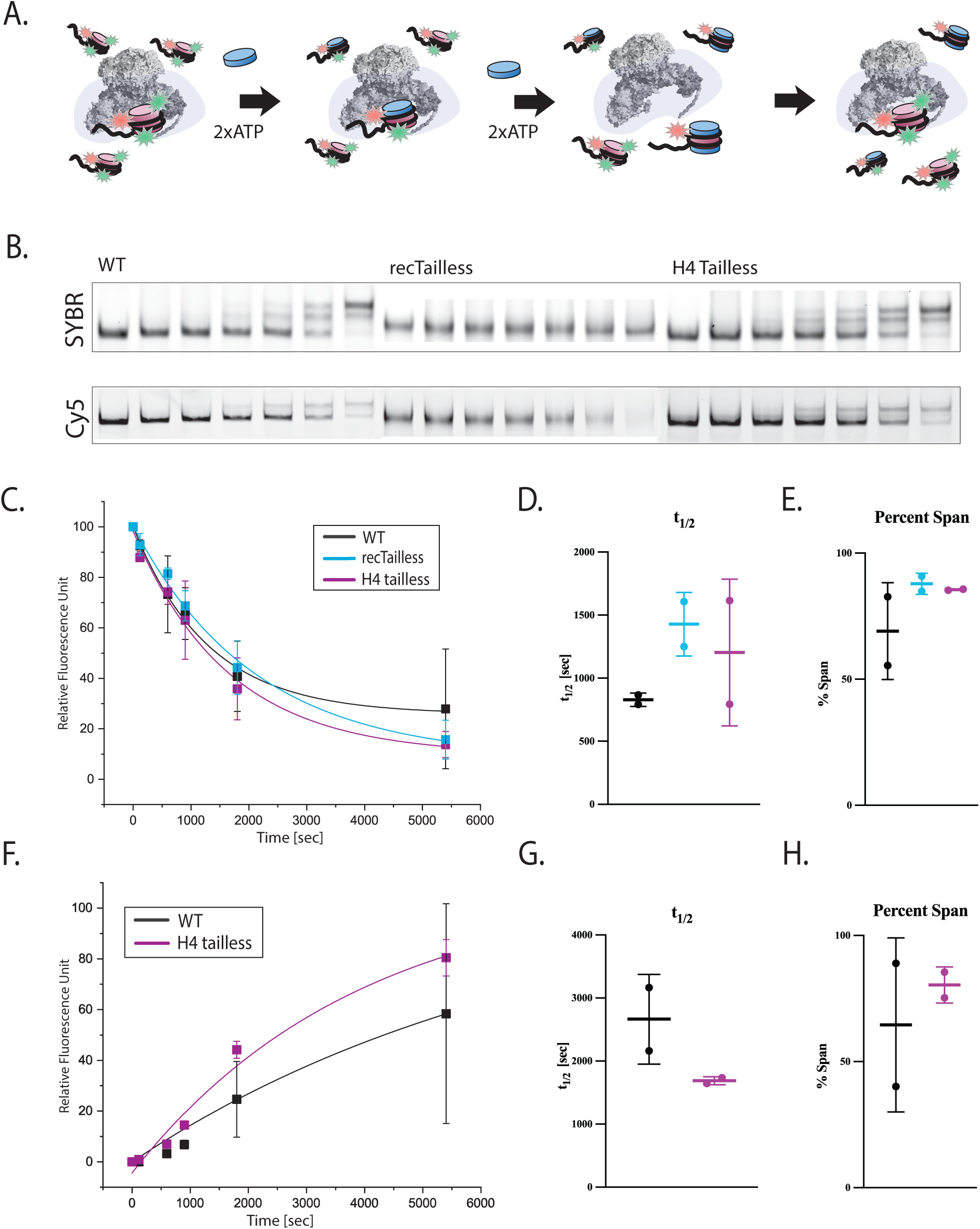
Histone tails do not impair SWR1C activity under catalytic conditions. **A.** Schematic of the ensemble FRET-based eviction assay under catalytic (multiple-turnover) conditions. Nucleosomes contain Cy5-labeled H2A and Cy3-labeled DNA; progressive replacement of Cy5– H2A/H2B dimers decreases FRET while SWR1C can process multiple substrates. **B.** Representative gel-based dimer-exchange assay under catalytic conditions. Reactions were sampled at 0, 1, 5, 10, 30, and 90 min after ATP addition. SYBR Gold staining (top) and Cy5 fluorescence (bottom) report deposition and eviction dynamics, respectively. **C–H.** Quantification of eviction (Cy5) **(C-E)** and deposition (SYBR Gold) **(F-H)** as indicated (each condition *N* = 2–3). Species pooling and scaling were performed as described in Figure 1. Kinetic models were fit to unscaled data to obtain parameters reported as mean ± SD. Statistics: Welch’s ANOVA with Dunnett’s T3 multiple comparisons for panels **D, E**; Welch’s *t* test for panels **G, H**. All additional comparisons/statistics are reported in the statistics table (Supplemental Table 1).

Dimer exchange reactions were performed under single turnover conditions (excess SWR1C enzyme to nucleosomal substrate) and contained free H2A.Z/H2B (ZB) dimers which act as an essential co-substrate (Luk et al., 2010; Singh et al., 2019) Addition of SWR1C to a wildtype nucleosome led to a rapid drop in FRET, showing the biphasic kinetics consistent with the sequential exchange of the two nucleosomal H2A/H2B (AB) dimers (Figure 1B) (Luk et al., 2010; Singh et al., 2021). Incubation of SWR1C with a nucleosome that lacks the N-terminal histone tails (recTailless) or the tail of histone H4 did not present any significant changes in the rates of either the initial, fast phase or the second, slow phase of the reaction (Figure 1B, 1C). An independent, deposition assay with recTailless and H4 tailless substrates surprisingly resulted in similar rates of fast and slow phases of the reaction (Supplemental Table 1A, 1B). In contrast, the fraction of reaction proceeding through the fast kinetic component differed significantly among conditions (percent fast phase, *p*=0.0167) (Figure 1D, Supplemental Table 1C). Dunnett’s T3 multiple comparisons indicated significant differences for WT vs RecTailless (*p*=0.0380) and for RecTailless vs H4Tailless (*p*=0.0477), while WT vs H4Tailless was not significant (*p*=0.8920) (Supplemental Table 1C). Together, these data indicate that tail removal does not substantially alter fitted half-lives but can shift amplitude ratios (Figure 1C-D).

As an independent approach, we employed a gel-based assay that monitors H2A.Z deposition (Luk et al., 2010). In this assay, nucleosome substrates were incubated with excess SWR1C, ATP, and free H2A.Z/H2B dimers in which H2B contains a 3xFLAG tag at its C-terminus. Reaction products are separated on native PAGE, and formation of the heterotypic (AB/ZB) and homotypic (ZB/ZB) nucleosomal products is detected by their reduced gel migration due to the 3xFLAG tag on H2B, as visualized by SYBR staining. These assays used nucleosomes that also harbor a Cy5-labeled H2A/H2B dimer, allowing the exchange reaction to also be followed by monitoring the loss of Cy5 fluorescence. In these reactions, nucleosomes were reconstituted with recombinant wild-type histone octamers, recombinant tail-less histone octamers (recTailless), or recombinant histone octamers that only lacked the H4 tail. Similar to the FRET-based assay, SWR1C catalyzed the stepwise deposition of H2A.Z. Exchange progressed to near completion by 60 min (Figure 1E). Surprisingly, nucleosomes that lack all histone tails (recTailless) did not show a change in mobility throughout the reaction progress in this gel-based assay, suggesting that such changes in mobility are not due solely to the increased mass imparted by the 3xFLAG epitope tag (Figure 1E), raising the possibility of altered electrophoretic behavior and/or altered product composition. In contrast, the loss of Cy5 signal supported eviction for recTailless substrates (Figure 1F). Quantification of gel-based deposition kinetics revealed a significant overall effect for the slow half-life (Welch’s ANOVA, p=0.0089), although corrected pairwise comparisons did not reach significance (e.g., WT vs RecTailless, p=0.0588) (Figure 1G, Supplemental Table 1C). Percent fast phase was not significantly different in this dataset (p=0.1268) (Figure 1H, 1K). In this set of gel-based dimer exchange reactions, fast-phase kinetics could not be determined during analysis.

Under catalytic (multiple-turnover) conditions, eviction and deposition time courses were best described by one-phase models within the sampled time. Across WT, RecTailless, and H4Tailless substrates, neither half-life nor percent span differed significantly (half-life, p=0.2679; percent span, p=0.6206) (Supplemental Table 1C). Together, these results indicate that unmodified histone tails are dispensable for robust SWR1C-mediated exchange in vitro and that the most reproducible tail-dependent effect is a shift in kinetic phase partitioning (% fast) rather than large changes in fitted half-lives.

### Extranucleosomal linker DNA strongly enhances SWR1C activity

Structural and biochemical studies have shown that several chromatin remodeling complexes interact with linker DNA to facilitate their catalytic activity, and some enzymes possess modules that function as rulers, capable of sensing the length of the flanking extra-nucleosomal DNA (Stokes and Perry, 1995; Saha et al., 2002; Stockdale et al., 2006; Yang et al., 2006; Farnung et al., 2017; Knoll et al., 2018; Zhou et al., 2018; Li et al., 2024). There is strong evidence that SWR1C also detects the length of the linker DNA, as its affinity to nucleosomes increases with increased linker DNA length (Ranjan et al., 2013; Poyton et al., 2022). This behavior is supported by SWR1C preferentially targeting the +1 nucleosomes flanked with longer nucleosome-free regions (Yen et al., 2013; Bagchi et al., 2019). Whether linker DNA enhances the catalytic activity of SWR1C subsequent to nucleosome binding remains unclear.

In order to assess the role of linker DNA on the activity of SWR1C, we prepared two types of nucleosomal substrates for use in the FRET-based, dimer exchange assay -- one lacking linker DNA (nucleosomal core particle, CP) and another, asymmetrical nucleosomal substrate featuring a 77 bp linker DNA on one side of the core particle. Both substrates were labeled with Cy3 on the linker-distal DNA end and with Cy5 on the histone H2A C-terminal tails. We tested varying concentrations (e.g., 10 nM, 20 nM, 30 nM, 60 nM) of SWR1C with constant substrate concentrations (Figure 3A). We hypothesized that if linker DNA primarily enhances the affinity of SWR1C for the nucleosome without influencing catalysis, the dimer exchange rates for both substrates would converge at higher enzyme concentrations, denoting substrate saturation. Interestingly, the deposition rates for both the core particle and the 77N0 nucleosome plateaued at ∼30-60 nM SWR1C (Figures 3A and 3B), but CP substrates exchanged more slowly than 77N0 substrates at matched enzyme concentrations. Pairwise Welch’s t tests comparing half-lives between nucleosome and CP conditions were significant at 30 nM SWR1C (p=0.0161) and 60 nM SWR1C (p=0.0125), with a similar trend at 20 nM (p=0.0566) (Supplemental Table 2C). These data indicate that linker DNA stimulates SWR1C activity independent of the differences in binding affinity.

**Figure 3.**
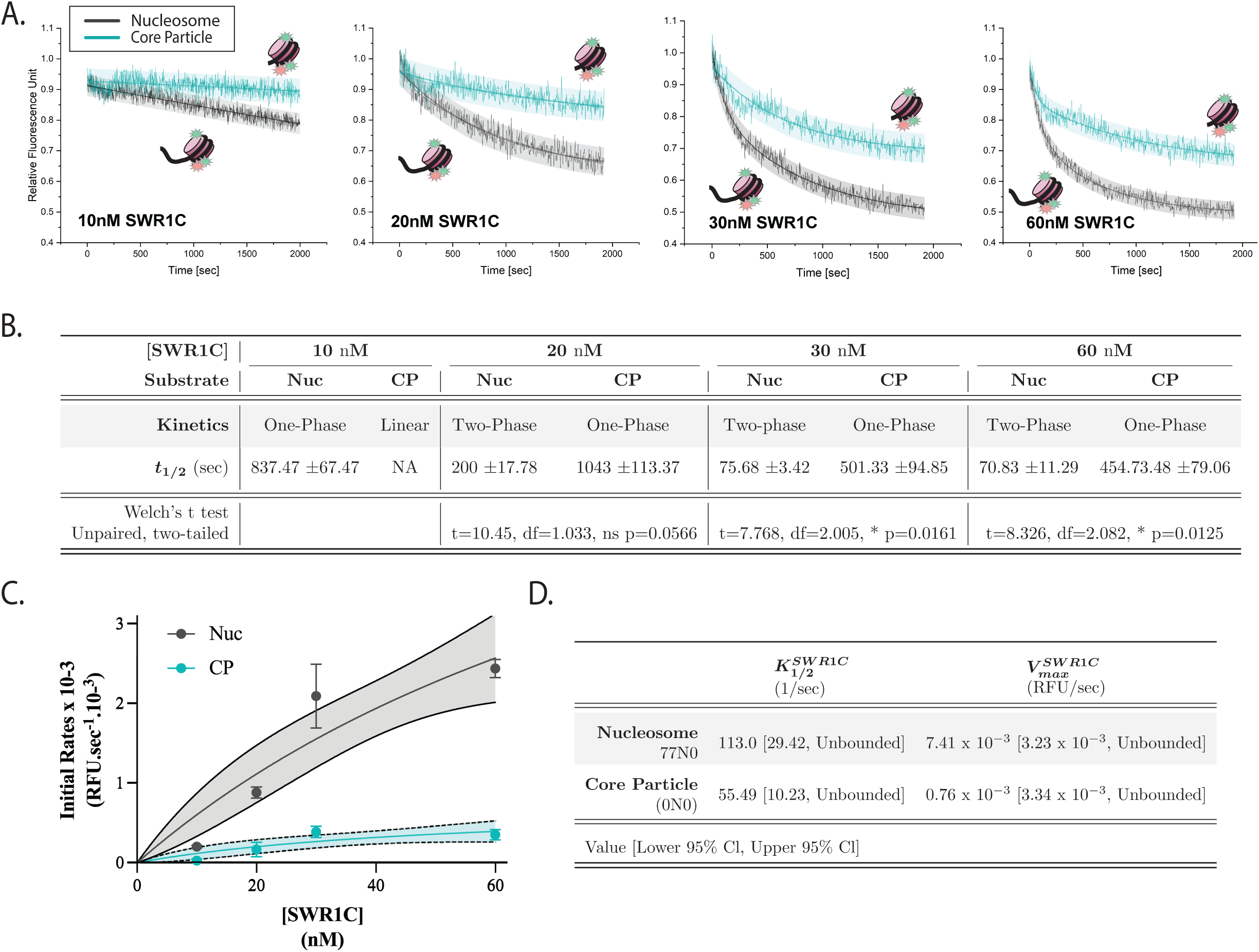
Linker DNA enhances SWR1C activity. **A.** Ensemble FRET-based eviction kinetics for nucleosomes versus core particles across SWR1C concentrations (10, 20, 30, and 60 nM; *N* = 2-3). Traces were normalized to their respective initial signals. **B.** Kinetic models (one-phase, two-phase, or linear as indicated) were fit to unnormalized data; fitted parameters are reported as mean ± SD. Shaded regions denote 95% confidence intervals of the fit. Welch’s *t* test is applied to the results. In comparison of kinetics data of 20 nM SWR1C with nucleosomes and core particles, W(t, df) = (10.45, 1.033) and p value = 0.0566. In comparison of kinetics data of 30 nM SWR1C with nucleosomes and core particles, W(t, df) = (7.768, 2.005) and p value = 0.0161. Finally, in comparison of kinetics data of 60 nM SWR1C with nucleosomes and core particles, W(t, df) = (8.326, 2.082) and p value = 0.0125. **C.** Initial rates (v_0_) derived from ensemble FRET-based dimer-exchange time courses for nucleosomes and core particles across SWR1C concentrations. For each replicate, the full time-course was fit using the best-supported kinetic model (one-phase or two-phase association/decay, as indicated for the corresponding dataset). The initial rate was then calculated as the instantaneous slope at reaction onset from the fitted model, 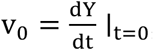 using the unscaled fit parameters. Initial rates are reported as mean ± SD across independent experiments (*N* = 2–3, as indicated). **D.** Initial rates were plotted as a function of SWR1C concentration and fit with a tQSSA rate equation to estimate apparent kinetic parameters (Supplemental Table 2A), apparent K_M_ and k_cat_ (*N* = 3). Fit confidence intervals are shown as indicated on the panel. Details can be found in Supplemental Table 2 (refer to Supplemental Table 2C for statistics).

To summarize early-time catalytic output across conditions, initial velocities (𝑣_0_) were estimated from fitted progress curves as the instantaneous slope at reaction onset (𝑡 = 0) (Figure 3C; Supplemental Table 2A). Initial velocities were then analyzed as a function of SWR1C concentration (Figure 3D). The linker-containing nucleosomes displayed substantially higher initial velocities than core particles across enzyme concentrations, consistent with linker DNA acting as a dominant substrate feature that promotes efficient exchange.

Kinetic parameters (𝐾_𝑀_) and (𝑘_𝑐𝑎𝑡_) for SWR1C-catalyzed histone exchange were determined by fitting initial velocities to the total quasi-steady state approximation (tQSSA), ensuring accurate modeling in the high-enzyme regime where substrate depletion by complex formation is non-negligible (Borghans et al., 1996; Tzafriri and Edelman, 2004; Pedersen et al., 2008) (Figure 3C-D, Supplemental Table 2A). Due to the lack of saturation observed in the enzyme kinetics data even at the highest feasible substrate concentrations, the individual parameters 𝐾_𝑀_and 𝑘_𝑐𝑎𝑡_could not be precisely determined and exhibited wide confidence intervals (as indicated by ’unconstrained’ 95% confidence interval, CI, numbers). Consequently, we focused our analysis on the specificity constant (𝑘_𝑐𝑎𝑡_⁄𝐾_𝑀_), which represents the catalytic efficiency under non-saturating conditions and is derived directly from the robust, linear slope of the initial velocity plots (Supplemental Table 2A). Represented by the specificity constant differences, the enzyme is approximately 4.5 times more efficient at processing the nucleosome than the core particle (Figure 3D).

### Dinucleosome context modestly alters phase partitioning without strong linker-length dependence

The majority of remodeler ATPases interact with nucleosomal DNA at superhelical location +/-2 (SHL 2), which is located adjacent to where the H4 tail domain protrudes from the nucleosome surface (Luger and Richmond, 1998; Mueller-Planitz et al., 2013; Racki et al., 2014). Indeed, biochemical and structural data have shown that several remodelers make direct contact with the H4 tail, supporting a direct mode of regulation on mononucleosome substrates. The H4 tail can bridge two nucleosomes by using a conserved basic patch to contact an acidic patch on an adjacent H2A/H2B dimer, an interaction that is believed to contribute to nucleosome-nucleosome interactions that facilitate chromatin higher-order folding (Fletcher and Hansen, 1995; Dorigo et al., 2003, 2004; Sinha and Shogren-Knaak, 2010; Pepenella et al., 2014; Chen et al., 2017). To test if SWR1C activity might be influenced by such nucleosome-nucleosome interactions, we reconstituted two dinucleosome substrates, where positioned nucleosomes were separated by either 30bp or 77bp of linker DNA. Each dinucleosome contained a 77bp nucleosome-free region at the 5’ end, and only the 3’ nucleosomal end was labeled with Cy3 (77N30N-Cy3, 77N77N-Cy3; Figure 4A). Although both nucleosomes harbor histone H2A labeled with Cy5, a FRET-based dimer exchange reaction primarily monitors the eviction of H2A from the distal nucleosome of the array.

**Figure 4.**
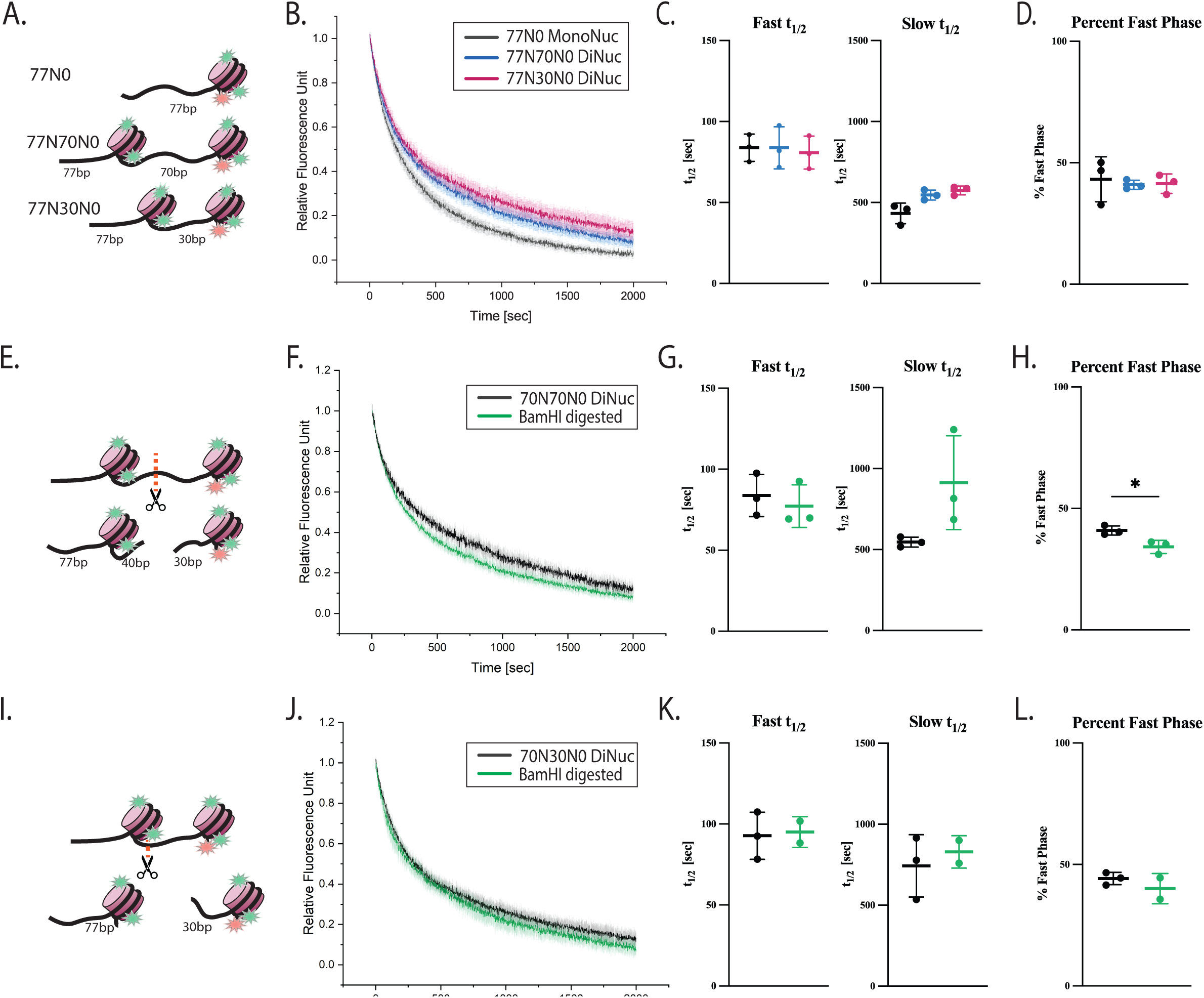
Mononucleosomes and dinucleosomes are processed similarly by SWR1C. **A.** Schematic of mononucleosome and dinucleosome substrates. All substrates contain Cy5-labeled H2A and Cy3-labeled DNA. Dinucleosomes contain either a long (70 bp) or short (30 bp) internucleosomal linker (77N70N0 and 77N30N0). The mononucleosome control (77N0) contains a 77-bp linker on one side and no linker on the other. **B-D.** Ensemble FRET-based eviction kinetics comparing SWR1C activity on mononucleosomes versus dinucleosomes (*N* = 3–4). **E, I.** BamHI digestion strategy to generate mononucleosome products from dinucleosomes. Digestion produces a FRET-invisible mononucleosome lacking Cy3 and a FRET-competent mononucleosome bearing a 30-bp linker (30N0), as diagrammed. **F, J.** Eviction kinetics for undigested versus BamHI-digested 77N70N0 (**F**) and 77N30N0 (**J**) substrates (*N* = 3). **G, H, K, L.** Quantification comparing dinucleosomes and BamHI-derived products (77N70N0 series: **G–H**, *N* = 3; 77N30N0 series: **K–L**, *N* = 2–3). Traces were scaled from basal to plateau (0–1) for visualization; kinetic fits were performed on unscaled data. Parameters are reported as mean ± SD. Shaded regions indicate 95% confidence intervals. Statistics: Welch’s ANOVA with Dunnett’s T3 multiple comparisons for panels **C, D**; Welch’s *t* test for **G, H, K, and L**. Parameters are reported as mean ± SD. Shaded regions indicate 95% confidence intervals. Detailed kinetics and statistics are reported in Supplemental Table 3.

To test how nucleosome context influences SWR1C exchange, we compared a mononucleosome substrate to dinucleosome substrates containing either a 70-bp or 30-bp internucleosomal linker (Figure 4A–D). Under single-turnover conditions, Welch’s ANOVA did not detect significant differences among mononucleosomes and dinucleosomes for fast half-life (p=0.9285), slow half-life (p=0.0769), or percent fast phase (p=0.9269) (Supplemental Table 3C). Thus, within this assay regime, internucleosomal linker length (30 vs 70 bp) does not measurably change fitted half-lives.

To decouple effects of array context from DNA template architecture, dinucleosome substrates were converted into mononucleosome products using a BamHI site within the internucleosomal linker (Figure 4E–L). BamHI digestion did not significantly alter fast or slow half-lives for either substrate. However, percent fast phase differed significantly for the 70-bp linker dinucleosome upon BamHI digestion (p=0.0292), whereas no significant change was detected for the 30-bp linker substrate (Supplemental Table 3C). These results suggest that nucleosome context can influence kinetic amplitude ratios in select configurations.

### Histone tail deletion increases the fast-phase contribution on dinucleosomes

To investigate whether the histone tail domains may have a larger regulatory role in the context of dinucleosomes, templates were reconstituted with recombinant tailless histone octamers (Figure 5A-F). For both long- and short-linker dinucleosomes, RecTailless substrates exhibited significant increases in percent fast phase relative to WT (70-bp linker, p=0.0481; 30-bp linker, p=0.0060), while differences in fast or slow half-lives did not reach significance (Supplemental Table 3C). Thus, as with mononucleosomes, tail removal primarily shifts the fraction of the reaction proceeding through the fast kinetic component. Notably, the increased rates were similar to those observed for mononucleosome substrates, indicating that unmodified histone tails may primarily regulate SWR1C at this primary level, rather than at the level of nucleosome-nucleosome interactions (Figure 5A-D).

**Figure 5.**
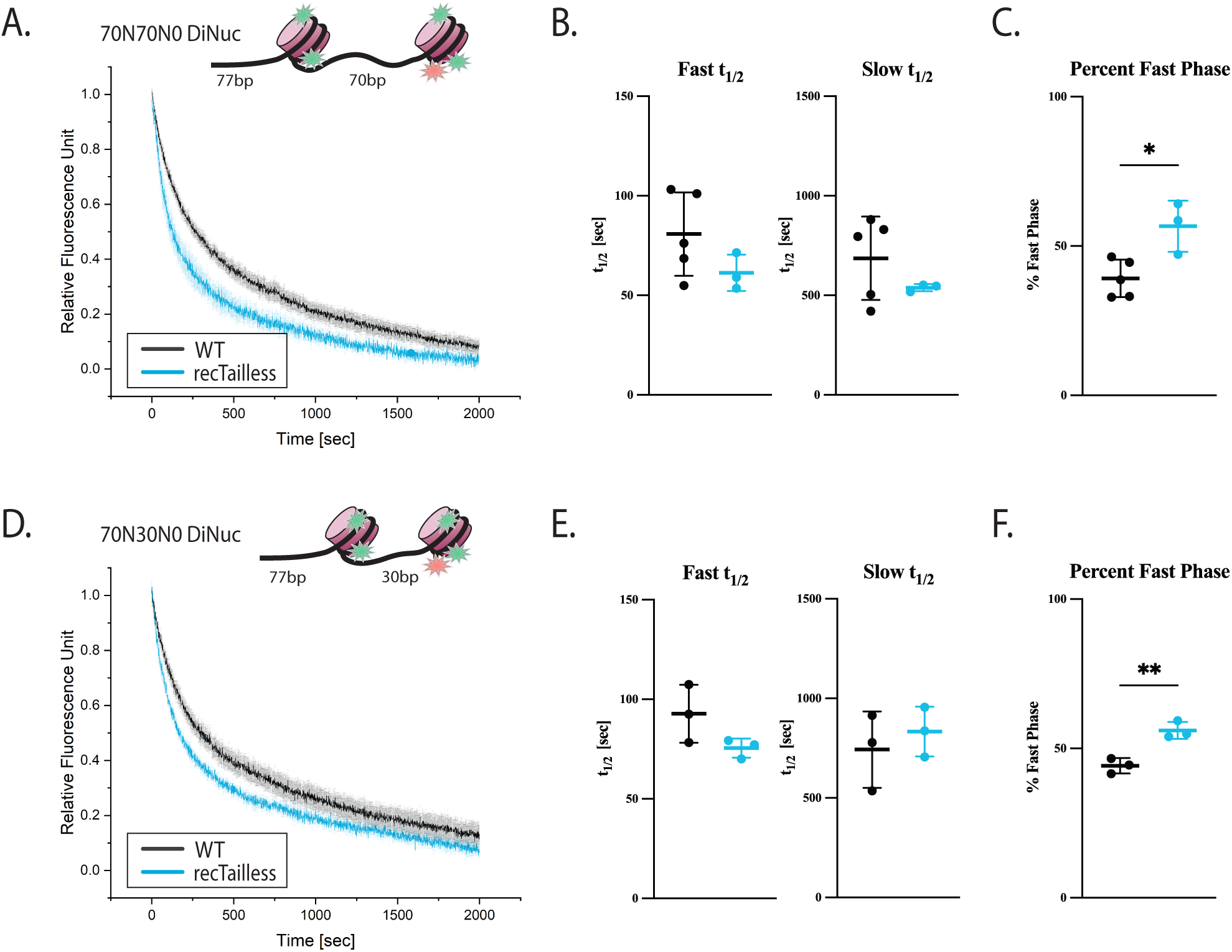
Histone tails modestly slow SWR1C-catalyzed exchange on dinucleosomes. **A-C.** Eviction kinetics on dinucleosomes with a long internucleosomal linker (70 bp) comparing WT and recombinant tailless substrates (*N* = 3). **D-F.** Eviction kinetics on dinucleosomes with a short internucleosomal linker (30 bp) comparing WT and recombinant tailless substrates (*N* = 2–3). Traces were scaled from basal to plateau (0–1) for visualization; kinetic fits were performed on unscaled data. Parameters are reported as mean ± SD; shaded regions indicate 95% confidence intervals. Statistics: Welch’s *t* test. Percent fast phase: long linker (panel C) t = 3.062, df = 3.317, p = 0.0481; short linker (panel F) t = 5.376, df = 3.959, p = 0.0060. Additional test statistics and pairwise comparisons, together with kinetic fit results, are reported in the Supplemental Table 3.

### Acetylation of histone tails facilitates H2A.Z deposition

Previous *in vivo* and *in vitro* studies have shown that acetylation of the lysine residues on H2A and/or H4 tails can stimulate H2A.Z deposition (Shia et al., 2006; Altaf et al., 2010; Ranjan et al., 2013; Cheng et al., 2015). The simplest model is that histone acetylation enhances the binding of SWR1C to nucleosomes, as the Bdf1 subunit contains two bromodomains (Ladurner et al., 2003; Matangkasombut and Buratowski, 2003; Zhang et al., 2005). However, one *in vitro* study that employed short, nucleosomal arrays suggested that histone acetylation by the NuA4 acetyltransferase may enhance SWR1C deposition activity, independent of an increase in nucleosome affinity. To investigate this further, we acetylated histones H2A and H4 with the NuA4 acetytransferase and then reconstituted both mononucleosome and dinucleosome substrates (Figure 6). We compared untreated nucleosomes (Nuc), NuA4-treated nucleosomes without acetyl-CoA (Nuc+NuA4), and NuA4-treated nucleosomes with acetyl-CoA (Nuc+NuA4+AcCoA) (Fig. 6A-F). Furthermore, mononucleosome assays for SWR1C activity were performed under both single turnover (Figure 6A-C) and multiple turnover conditions (Figure 6D-F). Under single-turnover conditions, Welch’s ANOVA detected no significant differences in fast half-life, slow half-life, or percent fast phase (Supplemental Table 4C), indicating that NuA4 acetylation does not play a direct role in SWR1C- mediated remodeling.

**Figure 6.**
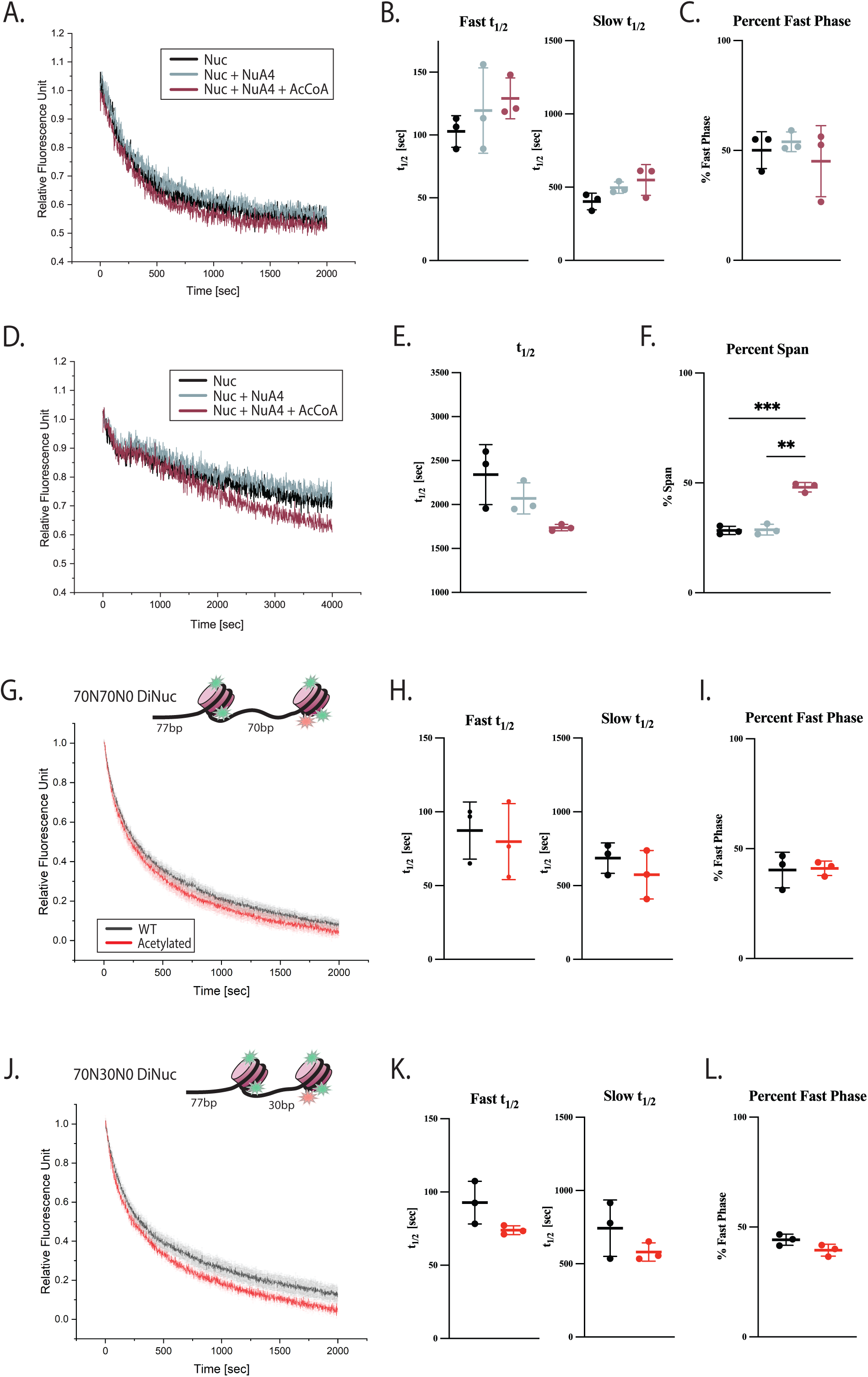
NuA4-mediated acetylation of histone tails enhances SWR1C activity under catalytic conditions. Ensemble FRET-based eviction assays performed on nucleosomes pretreated as indicated: untreated (Nuc), NuA4-treated without acetyl-CoA (Nuc+NuA4), and NuA4-treated with acetyl-CoA (Nuc+NuA4+AcCoA; acetylated). **A-C.** Single-turnover conditions. **D-F.** Catalytic conditions (*N* = 3). Signals were normalized from basal to plateau for visualization; kinetic models were fit to unscaled data. Fitted parameters are reported as mean ± SD, with 95% confidence intervals shown as shaded regions. Statistics: Welch’s ANOVA with Dunnett’s T3 multiple comparisons. Half-life: W(DFn, DFd) = 5.497 (2.000, 3.067), p = 0.0968. Reaction span/extent: W(DFn, DFd) = 70.66 (2.000, 3.951), p = 0.0008. Dunnett’s T3: Nuc vs Nuc+NuA4+AcCoA t = 11.88, df = 3.92, p = 0.0007; Nuc+NuA4 vs Nuc+NuA4+AcCoA t = 11.88, df = 3.92, p = 0.0007. **G-L.** Ensemble FRET-based eviction kinetics for acetylated dinucleosomes with **(A–C)** long and **(D–F)** short internucleosomal linkers (*N* = 3–5). Signals were normalized from basal to plateau for visualization; kinetic fits were performed on unscaled data and parameters are reported as mean ± SD, with 95% confidence intervals shown as shaded regions. Statistics: Welch’s *t* test. Additional test statistics and pairwise comparisons are reported in the Supplemental Table 4.

However, a reproducible, albeit small change in SWR1C activity was observed when mononucleosomes were in excess to SWR1C (multiple turnover conditions; Figure 6D-F). Under catalytic conditions, half-life differences were not significant (p=0.0968), whereas percent span differed strongly (p=0.0008). Dunnett’s T3 multiple comparisons indicated that acetylated nucleosomes differed from both untreated nucleosomes (p=0.0007) and NuA4-treated nucleosomes lacking acetyl-CoA (p=0.0013), while untreated nucleosomes and NuA4-treated nucleosomes lacking acetyl-CoA were not different (p=0.9954) (Supplemental Table 4C). These data indicate that NuA4-dependent acetylation increases the apparent extent of remodeling under enzyme-limiting conditions. These results would be consistent with a small impact of acetylation on enhancing increased substrate recruitment or binding stability where H4 tail acetylation serves as a docking signal.

The nucleosome-nucleosome interactions that are bridged by the H4 tail domain are disrupted by acetylation of lysine 16 of histone H4 (Tse et al., 1998; Shogren-Knaak et al., 2006). To test whether NuA4-mediated acetylation might stimulate SWR1C activity by such a mechanism, we compared SWR1C activity on dinucleosome substrates with and without NuA4 acetylation (Figure 6G-I and 6J-L). Acetylation of dinucleosome substrates under single-turnover conditions did not significantly alter fast half-life, slow half-life, or percent fast phase for either linker length (Supplemental Table 4C; Figure 6). Immunoblotting with a pan-acetyl-H4 antibody confirmed acetylation in NuA4-treated samples but also revealed cross-reactivity with acetylated H2A under these conditions (Supplemental Figure 4A).

## Discussion

ATP-dependent chromatin remodelers regulate chromatin accessibility and composition to support transcription, replication, and DNA repair (Clapier and Cairns, 2009; Clapier et al., 2017). SWR1C is unique among chromatin remodeling enzymes as it does not use the energy of ATP hydrolysis to mobilize or disrupt nucleosomes, but rather it edits nucleosome composition by exchanging H2A/H2B dimers with histone variant H2AZ/H2B dimers. Like the majority of remodeler ATPases, the Swr1 ATPase engages nucleosomal DNA at the internal SHL 2 position, placing it in close proximity to the protruding histone H4 N-terminal tail domain. The histone tails provide an important hub for protein-protein or protein-DNA interactions that regulate chromatin structure and dynamics, which can be further regulated through PTMs (Peng et al., 2021). Here, we used quantitative ensemble FRET- and gel-based assays to examine how histone tails, linker DNA, dinucleosome context, and NuA4-mediated acetylation influence SWR1C- catalyzed H2A.Z exchange. Three conclusions emerge. First, unmodified histone tails are largely dispensable for exchange kinetics in vitro, with the most consistent tail-dependent effect being a modest shift in phase partitioning (% fast) rather than robust changes in half-lives. Second, extranucleosomal linker DNA is a dominant determinant of efficient exchange, strongly enhancing activity relative to core particles lacking linker DNA. Third, dinucleosome context and NuA4 acetylation primarily modulate reaction partitioning/extent under specific regimes, with acetylation most clearly increasing reaction span under catalytic (enzyme-limiting) conditions.

### Histone tails minimally constrain SWR1C, primarily by altering phase distribution

Many remodelers engage nucleosomal DNA near SHL2 and are regulated by the adjacent H4 tail, including ISWI and CHD family enzymes for which H4 tail deletion strongly impairs activity (Clapier et al., 2001; Ferreira et al., 2007). Our data indicate that histone tails are not required for SWR1C function (Figure 1B), in contrast to remodelers, like ISWI and CHD1 (Clapier et al., 2001; Ferreira et al., 2007). In contrast, SWR1C-catalyzed exchange proceeded robustly on RecTailless and H4Tailless substrates, and fitted half-lives were not significantly altered in the single-turnover FRET assay (Figure 1 and Figure 2). Instead, we observed a significant change in the fraction of the reaction proceeding through the fast kinetic component (% fast) across tail conditions. This pattern argues against a strict mechanistic requirement for tails in the core exchange reaction and instead suggests that tails may modulate the distribution of kinetic pathways or substrate conformations that contribute to the ensemble progress curve. Importantly, because the assays report composite behavior of multi-step exchange, changes in Percent fast can reflect altered pathway weighting without requiring large shifts in intrinsic rate constants.

### Tailless substrates show altered electrophoretic behavior, limiting gel-based inference of deposition products

An interpretive challenge is that RecTailless substrates did not resolve the expected slower-migrating H2A.Z-containing species by SYBR Gold staining in the native gel assay, even though Cy5 signal loss supported eviction. Altered electrophoretic mobility of tailless nucleosomes (e.g., due to changes in charge distribution and/or DNA dynamics) could reduce resolution of heterotypic and homotypic products, complicating direct interpretation from migration alone. Alternatively, eviction without stable incorporation of H2A.Z/H2B cannot be excluded based solely on mobility shifts. One evidence that suggests SWR1C successfully completes H2A.Z deposition comes from FRET-based dimer exchange assay (Figure 2A, B). In this assay, Cy5 is tagged on the C-terminal tail of H2A.Z, leading to increased FRET activity as the H2A.Z/H2B dimer is deposited onto nucleosomes by SWR1C. Similar trends of fast and slow rates were observed under the dimer eviction assay (Figure 1B-D, 2A-B). Additionally, a direct test—such as anti-FLAG detection of gel-resolved products or H2A.Z-specific detection of incorporation—would resolve whether product composition differs for tailless substrates independent of mobility effects.

### Linker DNA is a dominant determinant of efficient SWR1C activity

In contrast to the modest effects of tail deletion, extranucleosomal linker DNA strongly enhanced SWR1C-mediated exchange. Core particles lacking linker DNA were significantly slower than 77N0 nucleosomes at matched SWR1C concentrations, including at enzyme concentrations where curves approach an apparent plateau. This supports a role for linker DNA in promoting productive engagement and/or catalytic steps downstream of binding. The biochemical requirement for linker DNA is consistent with SWR1C’s in vivo preference for depositing H2A.Z at nucleosomes flanking promoter nucleosome-free regions, where an adjacent stretch of nucleosome-free DNA is available.

To summarize the enzyme-titration analysis (Figure 3; Table 2), we fit progress-curve–derived onset velocities as a function of SWR1C concentration using a tQSSA-based framework to obtain apparent 𝑘_𝑐𝑎𝑡_, 𝐾_𝑀_, and 𝑘_𝑐𝑎𝑡_/𝐾_𝑀_values for nucleosome versus core-particle substrates. Because SWR1C-mediated exchange is multi-step/multi-substrate and our titration regime does not strictly satisfy classical steady-state assumptions, these parameters should be interpreted as effective descriptors of throughput and concentration dependence, with 𝑘_𝑐𝑎𝑡_/𝐾_𝑀_serving as a practical efficiency index for substrate comparison under matched conditions (Figure 3C-D; Supplemental Table 2A). Several 95% confidence intervals are reported as unconstrained (Supplemental Table 2A), reflecting limited parameter identifiability given the restricted enzyme range (10–60 nM) and higher variance at low enzyme concentrations (Supplemental Table 2B), which prevents the fit from bounding both the plateau and curvature required to define finite limits for all parameters. Unconstrained values are probably due to not reaching to the saturation concentrations of SWR1C. On the other hand, there is a distinct slope on both nucleosome and core particle substrates (Figure 3C), suggesting a clear comparable specificity constant, 𝑘_𝑐𝑎𝑡_/𝐾_𝑀_, values, which suggest a ∼4.45 fold efficiency for SWR1C when bound to nucleosomes with linker DNA (Figure 3D, Supplemental Table 2A).

Notably, the apparent tendency for core particles to exhibit a lower 𝐾_𝑀_-like concentration scale yet substantially lower catalytic output can be rationalized by kinetic trapping (Figure 7B). SWR1C may bind linkerless cores efficiently but populates nonproductive or weakly productive states that turnover slowly, whereas extranucleosomal linker DNA promotes productive engagement and ATPase coupling, thereby increasing effective catalytic throughput.

### Dinucleosome context modestly reshapes kinetics without strong internucleosomal linker-length dependence

Dinucleosome substrates were processed with broadly similar fitted half-lives across 30- and 70-bp internucleosomal linker lengths in our assay format, and omnibus statistical tests did not detect significant differences between mono- and dinucleosome substrates for the primary half-life metrics (Figure 4A-D). However, two observations indicate that array context can affect kinetic partitioning: (i) BamHI conversion of the 70-bp linker dinucleosome altered percent fast (Figure 8H), and (ii) RecTailless dinucleosomes displayed significantly higher percent fast relative to WT for both linker lengths (Figure 5C, F). Together, these findings are most consistent with the idea that nucleosome context influences the probability of entering a faster kinetic pathway (or the relative weighting of kinetic phases) rather than introducing a large barrier that uniformly changes fitted half-lives under the conditions tested.

### NuA4 acetylation increases reaction extent under catalytic conditions, consistent with enhanced recruitment or turnover

Regulation of chromatin remodeling complexes by histone acetylation has been well documented (Hamiche et al., 2001; Horn et al., 2002; Marilyn G. Pray-Grant et al., 2005; Shia et al., 2006; Altaf et al., 2010; Chatterjee et al., 2011, 2011, 2015; Ranjan et al., 2013; Sundaramoorthy et al., 2018). In the majority of cases, histone acetylation enhances the binding of a remodeler to its nucleosomal substrate, exploiting a bromodomain-containing subunit (Chatterjee et al., 2009, 2011). A previous study investigated whether acetylation of short, native yeast nucleosomal arrays stimulated H2A.Z deposition by SWR1C (Altaf et al., 2010). In this case, acetylation by the NuA4 acetyltransferase stimulated SWR1C activity, though it remained unclear if this stimulation was due solely to enhanced nucleosome binding affinity.

We performed a similar set of studies using our quantitative FRET-based dimer exchange assay and recombinant mononucleosomes and dinucleosomes (Figure 6). NuA4-mediated acetylation had minimal effects on fitted kinetic parameters under single-turnover conditions (Figure 6A-C), but it significantly increased reaction span under catalytic conditions (Figure 6D-F). This pattern is more consistent with enhanced productive engagement, recruitment, or turnover when enzyme is limiting than with acceleration of the intrinsic exchange step. Our assays may under-estimate the impact of histone acetylation, as our purification method leads to sub-stoichiometric levels of the Bdf1 subunit that harbors two bromodomains (Watanabe et al., 2015). Bdf1 is required for optimal recruitment of SWR1C to target loci *in vivo* (Ladurner et al., 2003; Zhang et al., 2005), but it is not required for H2A.Z deposition, *in vitro* (Wu et al., 2009). Notably, acetylation of dinucleosome substrates did not produce significant changes in half-lives or percent fast under single-turnover conditions (Figure 6G-L), suggesting that the dominant effect of acetylation in our system is conditional and regime-dependent. In contrast, our results indicate that SWR1C activity is strongly modulated by the length of the nucleosomal linker, both at the level of nucleosome binding affinity and catalytic activity. Regulation by adjacent free DNA is likely mediated by the Swc2 subunit, which may couple linker DNA sensing to the Swr1 ATPase (Girvan et al., 2024).

Key limitations include the altered mobility of tailless substrates in native gels (necessitating direct verification of H2A.Z incorporation), limited enzyme-titration coverage above 60 nM for constraining half-maximal parameters, and incomplete quantification of acetylation stoichiometry due to antibody cross-reactivity. Future experiments that directly detect incorporation products (anti-FLAG or H2A.Z-specific detection), quantify acetylation levels more precisely, and measure SWR1C binding/ATPase coupling as a function of linker DNA would clarify which mechanistic step(s) are most sensitive to these regulatory cues.

## Materials and Methods

### SWR1C Isolation

SWR1C was isolated from yeast as previously described (Mizuguchi et al., 2004; Singh et al., 2019; Gutierrez et al., 2022). Briefly, the yeast strain expressing C-terminally 3xFLAG-tagged Swr1 in an Δhtz1 background (strain yEL90, a gift from Dr. Ed Luk, SUNY Stony Brook) was grown in enriched media YPD or in enriched media with supplemented with 50mg/L adenine YPAD to an OD of ∼3 (YPD) or ∼6 (YPAD). Cells were collected by centrifugation, resuspended in minimal volume, frozen as pebbles/noodles, and stored in -80 °C. Yeast pebbles were cryogenically ground using a planetary ball cryomill (Retsch PM100) until a fine powder was reached.

All buffers contained freshly added: DTT, PMSF, Pepstatin A, Leupeptin, Benzamidine, Chymostatin, and Aprotinin. Powdered yeast was resuspended (1:1 w/v) in lysis buffer (25mM HEPES, 1mM EDTA, 300mM KCl, 10% glycerol, 0.02% NP-40, 10mM b-glycerophosphate, 0.5mM NaF, 1mM Na-butyrate, pH to 7.6 with KOH) and incubated at room temperature until homogeneous, followed by 30 min incubation at 4 °C. Lysates were clarified by centrifugation at 35,000 rpm for 2h at 4 °C in a Ti-45 Beckman rotor.

The supernatant is transferred to a container without disturbing the pellet and was incubated rotating at 4°C for 3 h with 3xFLAG affinity resin (Sigma-Aldrich M2 resin) pre-equilibrated in lysis buffer (3 x 10 mL washes). The resin was pelleted by centrifugation at 1,000 rcf for 2 min at 4 °C, transferred to a gravity column, and washed 3 times with 25 mL of B-0.5 buffer (25mM HEPES, 1mM EDTA, 2mM MgCl_2_, 500mM KCl, 10% glycerol, 0.02% NP-40, 10mM β-glycerophosphate, 0.5mM NaF, 1mM Na-butyrate, pH to 7.6 with KOH). This was followed by 3 times washes with 10 mL of B-0.1 buffer (25mM HEPES, 1mM EDTA, 2mM MgCl_2_, 100mM KCl, 10% glycerol, 0.02% NP-40, 10mM b-glycerophosphate, 0.5mM NaF, 1mM Na-butyrate, pH to 7.6 with KOH).

SWR1C was eluted by rotating at 1 mL of B-0.1 buffer containing 0.5 mg/mL 3xFLAG peptide (Sigma, F4799) for 30 min at 4 °C. The eluate was clarified by centrifugation at 1,000 rcf for 2 min at 4 °C, and the supernatant was collected, making sure there is no contamination from the resin; a small amount was saved for quantification of the protein. The supernatant was aliquoted, flash frozen in liquid nitrogen, and stored at -80 °C. Protein concentration was measured by running the protein on SDS-Page gel and comparing the concentration to a BSA standard.

### NuA4 isolation

NuA4 histone acetyltransferase complex was isolated from *Saccharomyces cerevisiae* expressing TAP-tagged Epl1, as described previously (Manning and Peterson, 2014). Yeast pellets were prepared for protein purification as explained in SWR1C isolation. The cells are then ground using a Retsch cryomill. The powder was allowed to thaw on ice and resuspended in lysis buffer. The lysate is clarified via ultracentrifugation at 40,000 rpm for 1 h at 4 °C in a Ti-45 Beckman rotor, and the supernatant was incubated with IgG Sepharose resin for 3 h at 4 °C to bind the Protein A epitope.

The resin was washed extensively with lysis buffer to remove unbound proteins. The protein of interest was cleaved from the resin by incubation with TEV protease in TEV cleavage buffer overnight at 4 °C. The following day, the eluate was incubated with calmodulin resin equilibrated in calmodulin binding buffer. After washing, bound protein was eluted using calmodulin elution buffer.

The eluate was dialyzed overnight at 4 °C against dialysis buffer to remove EGTA and then concentrated to ∼150 μl. Protein was flash frozen in liquid nitrogen and stored at -80 °C.

The buffers used in this protocol include E Buffer (20 mM Hepes, pH 7.4, 350 mM NaCl, 0.1% Tween 20, 10% Glycerol), Lysis Buffer (E Buffer with protease inhibitors), TEV Cleavage Buffer (E Buffer with protease inhibitors and 1 mM DTT), Calmodulin Binding Buffer (E Buffer with 2 mM CaCl2, protease inhibitors, and 1 mM DTT), Calmodulin Elution Buffer (E Buffer with 10 mM EGTA, protease inhibitors, and 1 mM DTT), and Dialysis Buffer (E Buffer with PMSF and 1 mM DTT).

All buffers were freshly prepared, kept on ice or at 4 °C, and supplemented with protease inhibitors immediately prior to use. Buffer components were thoroughly dissolved to ensure homogeneity and avoid bubble formation.

### Histones expression constructs

In this study, yeast H2A, H2A.Z, and H2B, and Xenopus H3 and H4 were used to prepare hybrid nucleosomes. The plasmids containing histones H3 (Δ1-37) and H4 (Δ1-23) without histone tails were donated from Richmond lab. To investigate the role of histone tails in SWR1C function, N-terminal tails of histones H2A, and H2B were truncated using Gibson assembly, and verified by Sanger sequencing. H2A N-tail residues 1-13 and H2B N-tail residues 1-23 were deleted to remove the N-terminal tails.

### Nucleosome reconstitution

Mononucleosome substrates were assembled as either core particles (CP; 147 bp, no extranucleosomal linker DNA) or asymmetric linker nucleosomes (77N0; 147 bp core plus a 77-bp linker on one side; total 224 bp). Dinucleosome substrates were assembled on a 601–603 template containing a 77-bp 5′ nucleosome-free region followed by two positioned nucleosomes separated by either a 70-bp or 30-bp internucleosomal linker (77N70N0 and 77N30N0). In this nomenclature, ’77’ indicates the 5′ extranucleosomal DNA length, ’N’ denotes a positioned nucleosome, the middle number indicates internucleosomal linker length, and terminal ’0’ indicates no additional linker DNA at the 3′ end of the distal nucleosome.

Nucleosomes were reconstituted as previously described (Luger et al., 1999). Recombinant histones were expressed in *Escherichia coli* Rosetta 2 (DE3) cells with or without pLysS plasmid and purified under denaturing conditions. Histone dimers and octamers were assembled by stepwise dialysis and gel filtration. For fluorescence-based assays, cysteine residues were introduced via site-directed mutagenesis to allow site-specific labeling with Cy3 and Cy5 maleimide fluorophores.

DNA fragments containing 601 or 601-603 sequences with linker lengths of 30 or 70 bp were PCR-amplified from plasmids using unlabeled or 5’ end-labeled primers (IDT). PCR reactions were performed using 0.1ng/μL plasmid template, 500 nM each of forward and reverse primers (labeled or unlabeled depending on the experiment), 200 μM dNTPs, 0.02 U/μL Phusion DNA Polymerase (NEB), and 1x Phusion High-Fidelity Buffer.

Nucleosome reconstitution was carried out by salt-gradient dialysis. Mononucleosomes were assembled 1:1 molar ratio of DNA to histone octamer; dinucleosomes arrays used 1:2 DNA:octamer ratio. DNA and histones were mixed in 600 μL of octamer buffer (10 mM Tris at pH 7.4, 1 mM EDTA, 2M KCl, and 5 mM Beta-mercaptoethanol or 1 mM DTT). Dialysis was performed overnight at 4 °C into 3L of nucleosome buffer (3 liters of nucleosome buffer: 10 mM Tris at pH 7.4, 1 mM EDTA, 50 mM KCl, and 5 mM Beta-mercaptoethanol or 1 mM DTT).

Nucleosome assembly was verified by native polyacrylamide gel electrophoresis (native PAGE), and free DNA, mononucleosomes, or dinucleosomes compositions were assessed based on electrophoretic mobility.

### Histone Acetylation

Histone acetylation was carried out using the NuA4 histone acetyltransferase complex, following a modified protocol based on the Coté laboratory method (Altaf et al., 2010). Acetylation reactions were assembled in a final volume of 50 μL, containing 500 nM nucleosomes and 0.15 mM acetyl-CoA (Sigma-Aldrich) in nucleosome buffer (10 mM Tris at pH 7.4, 1 mM EDTA, 50 mM KCl, and fresh 5 mM Beta-mercaptoethanol or 1 mM DTT). Approximately 5 μL of concentrated NuA4 complex was added per reaction. The same batch of NuA4 was used for all experiments to ensure consistent enzymatic activity. Reactions were mixed gently and incubated at room temperature in the dark for 45 min.

Acetylation efficiency was assessed by SDS-PAGE and western blotting. Samples were resolved on an 18% SDS-PAGE gel, transferred to a PVDF membrane and probed with an anti-pan-acetyl histone H4 antibody (Histone H4ac (pan-acetyl) antibody, pAb, 39925, Active Motif) at 1:1000 dilution. Wildtype, nonacetylated nucleosomes were included as negative controls. Signal was detected using appropriate HRP-conjugated secondary antibodies and chemiluminescence reagents.

### FRET-based dimer exchange assays

FRET-based H2A.Z dimer exchange assays were performed as previously described (Singh et al., 2019). Final reaction concentrations were 12 nM nucleosomes, 70 nM H2A.Z/H2B dimers, and either 30 nM SWR1C (single-turnover conditions) or 2 nM SWR1C (catalytic conditions), in remodeling buffer (25 mM HEPES at pH 7.6, 0.2 mM EDTA, 5 mM MgCl_2_, 70 mM KCl, and fresh 1 mM DTT). SWR1C concentrations were doubled for di-nucleosome experiments to keep nucleosome:SWR1C ratio stable. Reactions were initiated by addition of 1 mM ATP, gently mixed by pipetting and immediately transferred to the spectrophotometer.

Fluorescence was recorded at 30 °C using either ISS PC1 spectrophotometer or Tecan Infinite M1000 PRO microplate reader (excitation: 530 nm for Cy3; emission: 670 nm for Cy5). Measurements were taken up to 2000 seconds / ∼ 30 min for single-turnover conditions or 4000 seconds / ∼ 60 min for catalytic experiments. Single-turnover conditions were defined as SWR1C in excess of nucleosome; catalytic conditions were defined as nucleosome in excess of SWR1C.

For enzyme-titration experiments, SWR1C was varied (10, 20, 30, 60 nM) at fixed nucleosome (12 nM), ZB dimer (70 nM), and ATP (1mM) concentrations; all other conditions were identical.

### Gel-based dimer exchange assay

Gel-based assays were carried out as previously described (Ranjan et al., 2013). A 6x master mix (130 μl total) was assembled using the same concentrations and buffer conditions as FRET-based dimer exchange assay. Reactions were initiated with 1 mM ATP and incubated at room temperature in the dark.

Aliquots were taken at 0, 1, 5, 10, 15, and 60 min (single-turnover), or 0, 1, 5, 10, 30, and 90 min (catalytic). Reactions were quenched with 1 μg plasmid DNA, incubated on ice in the dark, and resolved on 6% native PAGE. Gels were imaged for Cy3, Cy5, and SYBR Gold (ThermoFisher) fluorescence using a GE Typhoon scanner. Band intensities were quantified using FIJI (ImageJ) Gel Analysis.

For eviction quantification, ACy5/ACy5 and ACy5/Z species were pooled; for deposition quantification, ACy5/Z and Z/Z species were pooled.

### Data Analysis

All experiments were conducted with at least two replicates per condition. For plotting only, traces were normalized to the signal value at x=0 (Figure 6A) or scaled from basal to plateau (0–1) . All kinetic fitting and parameter extraction were performed on unscaled, unnormalized (raw) data. Gel band intensities were background-subtracted and normalized relative to total signal. Data were analyzed in GraphPad Prism (v10), OriginLab Pro and Python (scripts can be found in Appendix III, and fit to standard linear, one-phase, or two-phase decay models.

### Exponential model fitting and half-life calculation

Time courses were fit by nonlinear regression to one-phase or two-phase exponential association/decay models (as indicated per dataset). For one-phase exponentials, 𝑡_1/2_ = ln (2)/𝑘. For two-phase fits, fast- and slow-phase half-lives were computed from 𝑘_fast_and 𝑘_slow_; the percent fast phase was computed from the fitted amplitudes: %𝑓𝑎𝑠𝑡 = 100 ⋅ 𝐴_fast_/(𝐴_fast_ + 𝐴_slow_). Model selection was guided by residual analysis and parsimony; residual plots supporting model choice (Figure 3 and 5) and equation fit metrics (Table 1B, 2B, 3B, and 4B) are provided.

### Initial-rate determination

Initial rates (𝑣_0_) were derived via two different methodology and compared; through early linear window (data not shown) and fitted progress-curve models (Figure 7A, Table 2A) as the instantaneous slope at reaction onset (Cornish-Bowden, 1975; Duggleby, 1985; Bursch et al., 2023). The rates showed similar trends under both methods. The initial velocities obtained from progress curves presented in this work, because progress curves exhibited curvature and the initial linear regime can be short, initial velocities were estimated from fitted progress curves to reduce subjectivity in selecting an “early linear” window (Cornish-Bowden, 1975; Duggleby, 1985, 1995).

Initial velocities (𝑣_0_) were derived from the **first derivative** of the fitted progress curves at 𝑡 = 0.

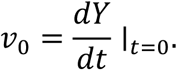

For multi-phasic eviction data, 𝑣_0_was calculated as the **sum of the weighted initial rates** of the individual phases (𝑣_0_ = ∑𝑘_𝑖_ ⋅ Span_𝑖_)."

For linear fits (using data normalized to initial signal: y, at x=0),

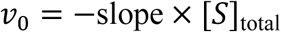

For one-phase exponentials,

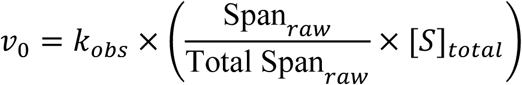

For two-phase decay fits,

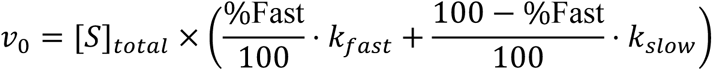

where [S] is nucleosome or core particle concentration. This approach avoids subjective selection of an “early linear window” and is consistent with established progress-curve analysis methods.

### Concentration dependence of initial rates

The dependence of the initial rate (𝑣_0_) on total enzyme concentration (𝐸_𝑇_) was modeled using the total Quasi-Steady State Approximation (tQSSA), which accounts for conditions where enzyme and substrate concentrations are comparable:

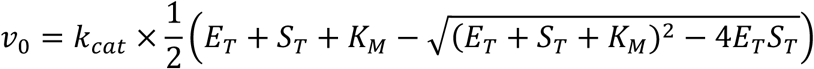

where 𝑆_𝑇_is the total nucleosome concentration, 𝑘_𝑐𝑎𝑡_is the catalytic turnover number, and 𝐾_𝑀_is the Michaelis constant (Borghans et al., 1996; Tzafriri and Edelman, 2004; Pedersen et al., 2008; Choi et al., 2017; Shin et al., 2024).

### Statistics

All statistical tests were performed using GraphPad Prism 10. For comparisons among ≥3 conditions, Welch’s ANOVA was used, followed by Dunnett’s T3 multiple comparisons. For two-condition comparisons, two-tailed unpaired Welch’s *t* tests were used. Exact test statistics, degrees of freedom, and *p* values are reported in the figure legends and the statistics tables (Supplemental Tables 1C, 2C, 3C, and 4C).

## Acknowledgements

We thank Timothy Richmond and Ed Luk for their bacterial and yeast strain contributions. We thank all the Peterson lab members for their feedback and assistance. This work was supported by the National Institutes of Health [R35-GM122519 to C.L.P.].

## Supplemental Figures

**Supplemental Figure 1.**
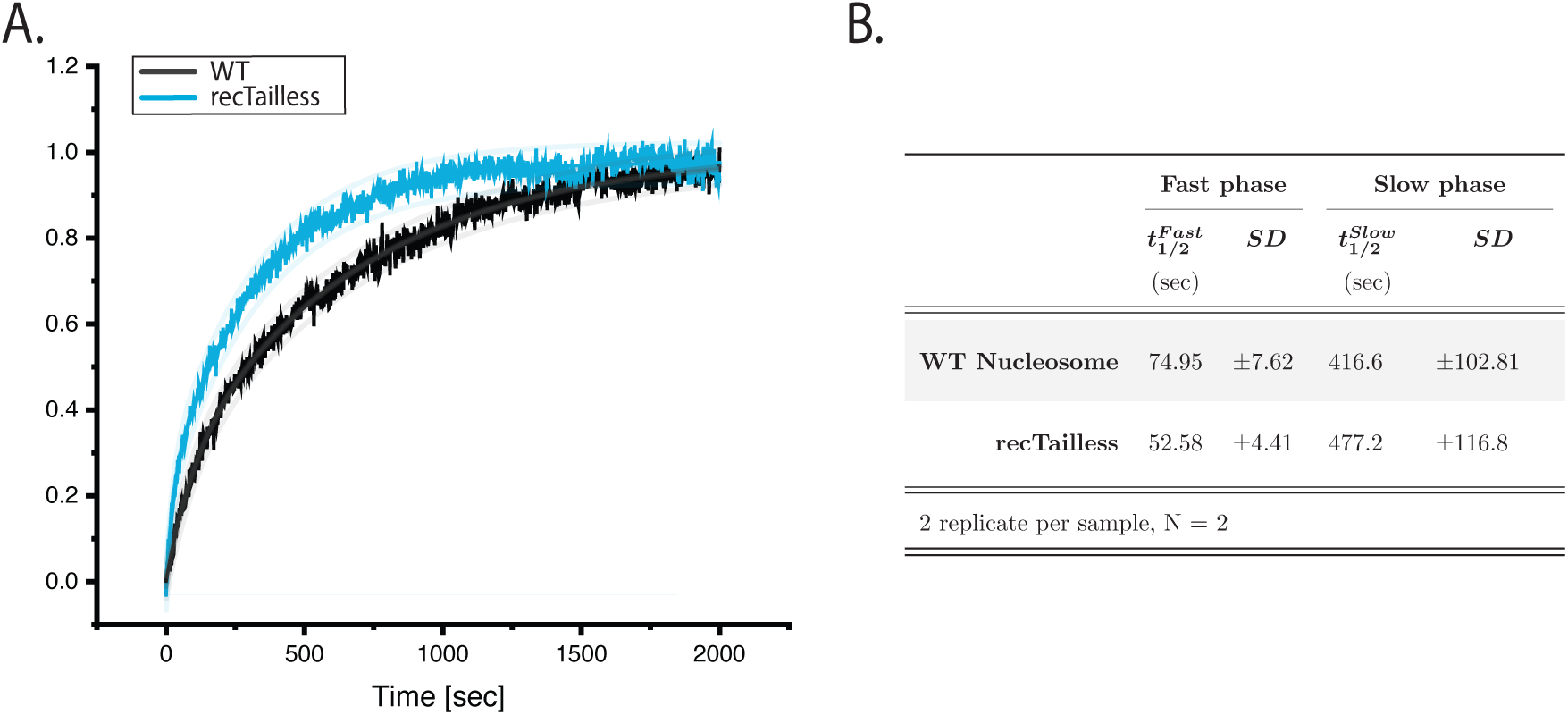
Deposition kinetics show trends similar to those of eviction kinetics. **A.** Ensemble FRET-based deposition kinetics comparing WT (black) and recombinant tailless (blue) nucleosomes (*N* = 3). **B.** Summary of fitted kinetic parameters (fast/slow t₁/₂ and fast-phase fraction) for traces in (A). Data were scaled from basal to plateau (0–1) for visualization; fits were performed on the unscaled data using one-phase or two-phase association/decay models as indicated. Parameters are reported as mean ± SD. Details of these reactions can be found in Supplemental Table 1.

**Supplemental Figure 2.**
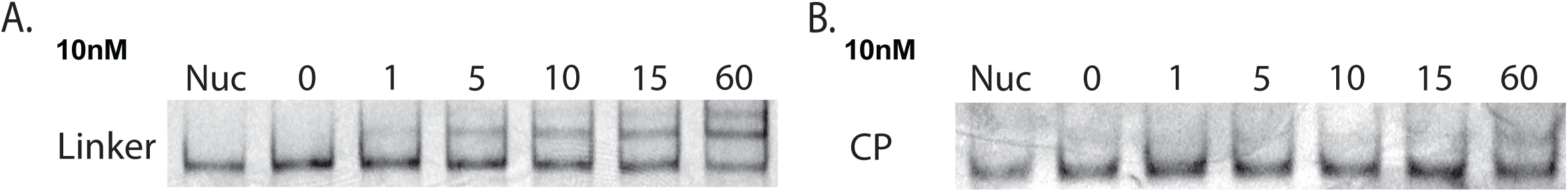
Representative images of gel-based dimer exchange with 10 nM SWR1C. **A-B.** Representative gel-based dimer-exchange assays at 10 nM SWR1C using nucleosomes (**A**) or core particles (**B**).

**Supplemental Figure 3.**
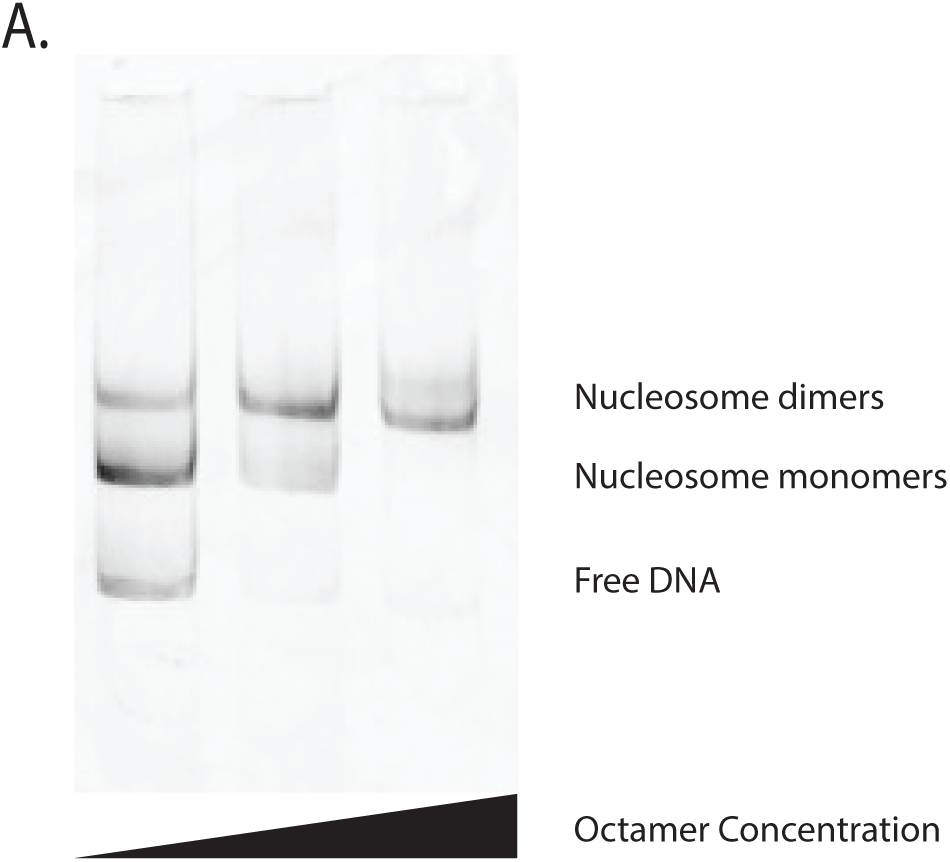
Reconstitution of nucleosome dimer across a range of octamer concentrations. **A.** Native PAGE of assembly reactions performed by salt dialysis on DNA template containing 601 and 603 positioning sequences. Bands corresponding to free DNA, mononucleosome, and dinucleosome are marked at right

**Supplemental Figure 4.**
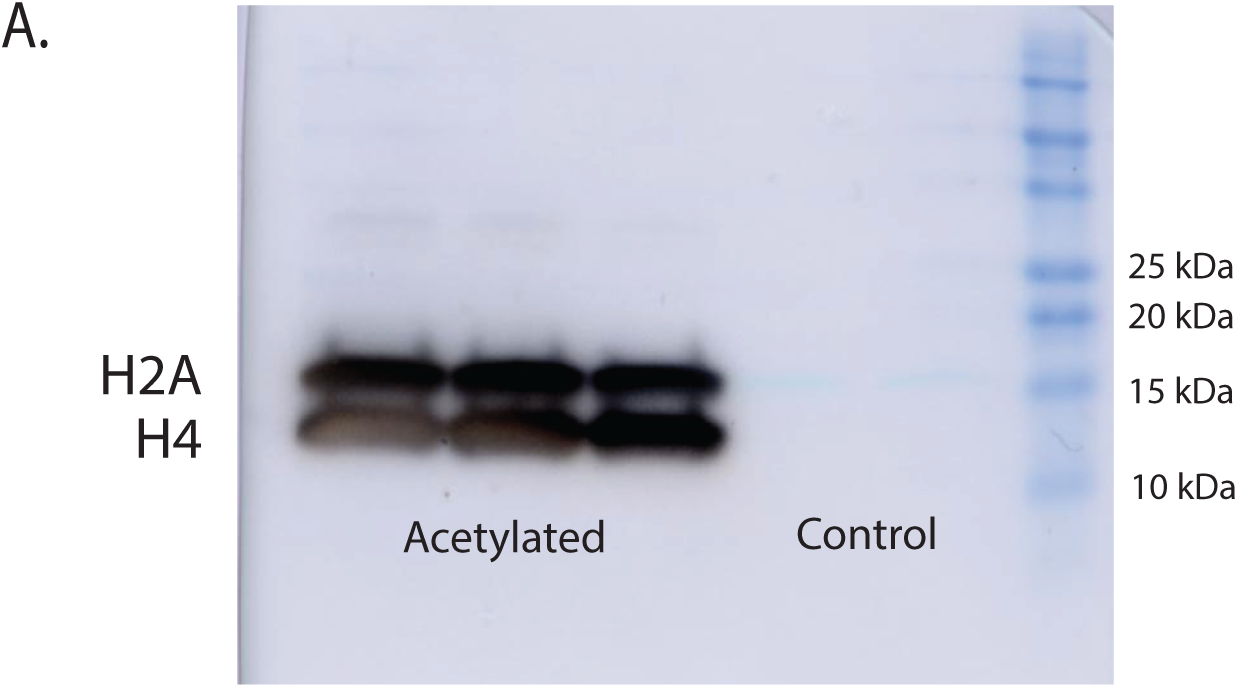
A pan-acetyl-H4 antibody detects NuA4-acetylated H4 and cross-reacts with acetylated H2A. **A.** Western blot of NuA4-treated nucleosomes probed with a pan-acetyl-H4 antibody. The antibody detects acetylated H4 and shows cross-reactivity with NuA4-acetylated H2A under these conditions (validated independently by fluorophore-tagged H2A signal; not shown).

**Supplemental Table 1.**
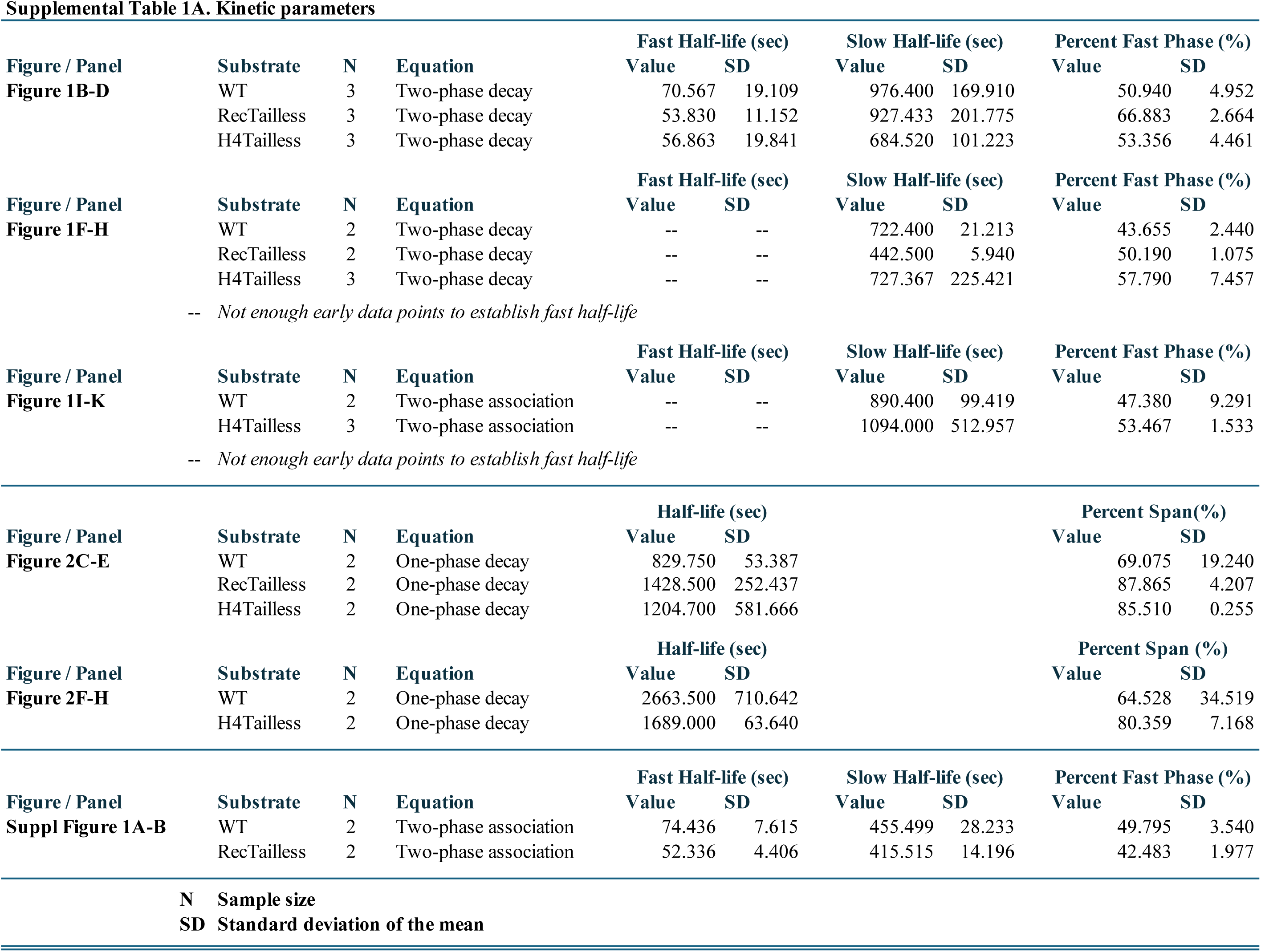
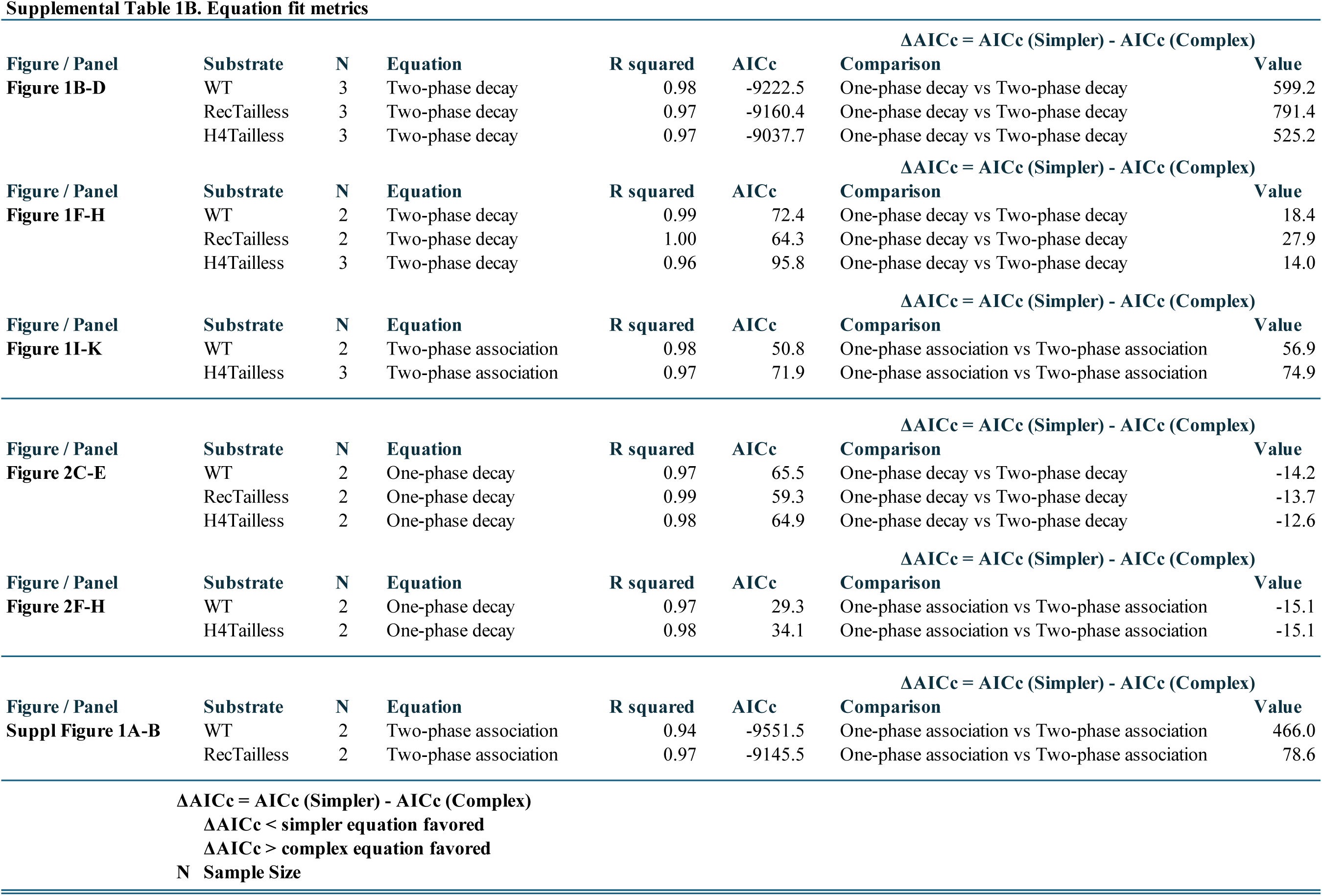
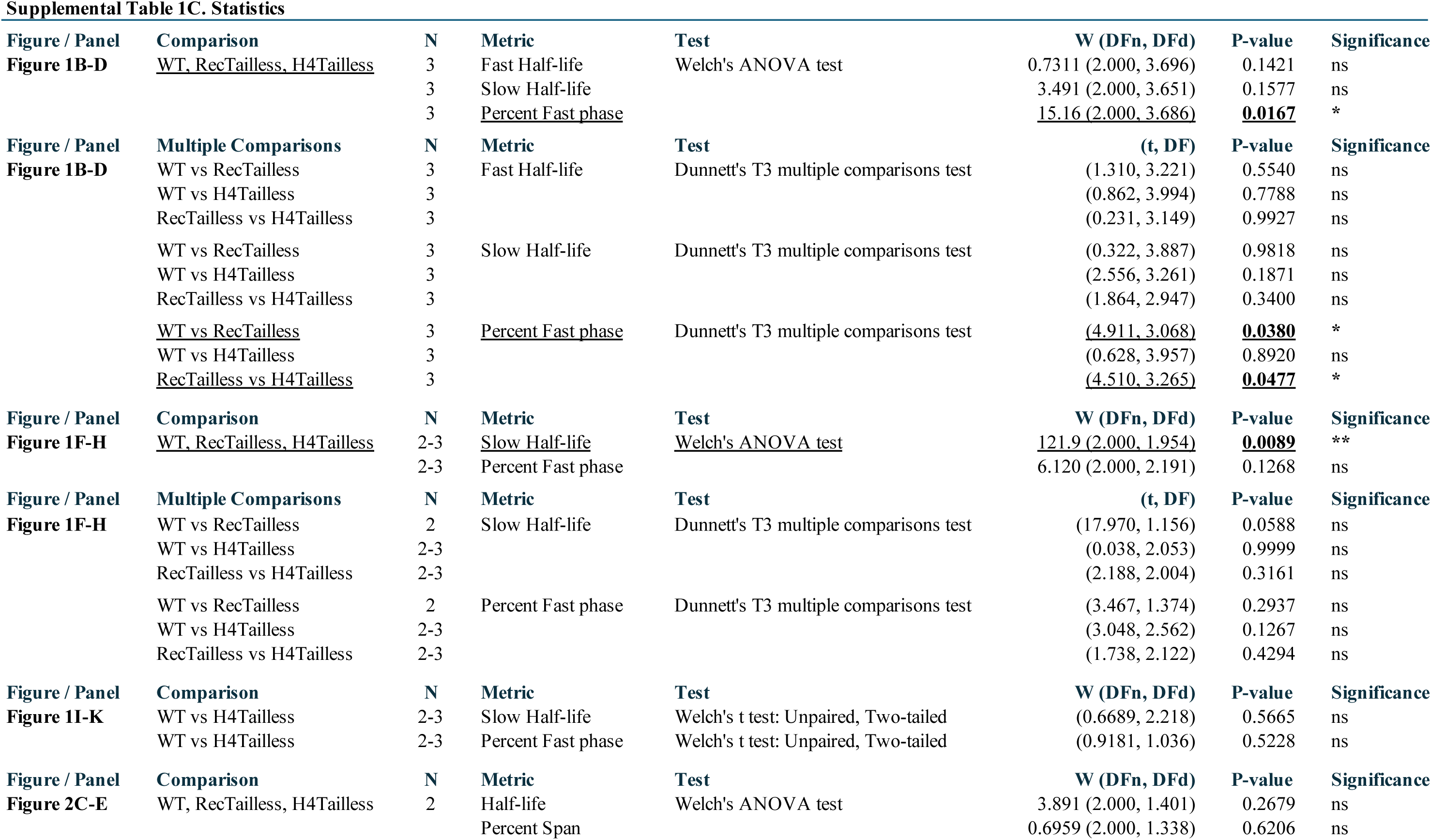
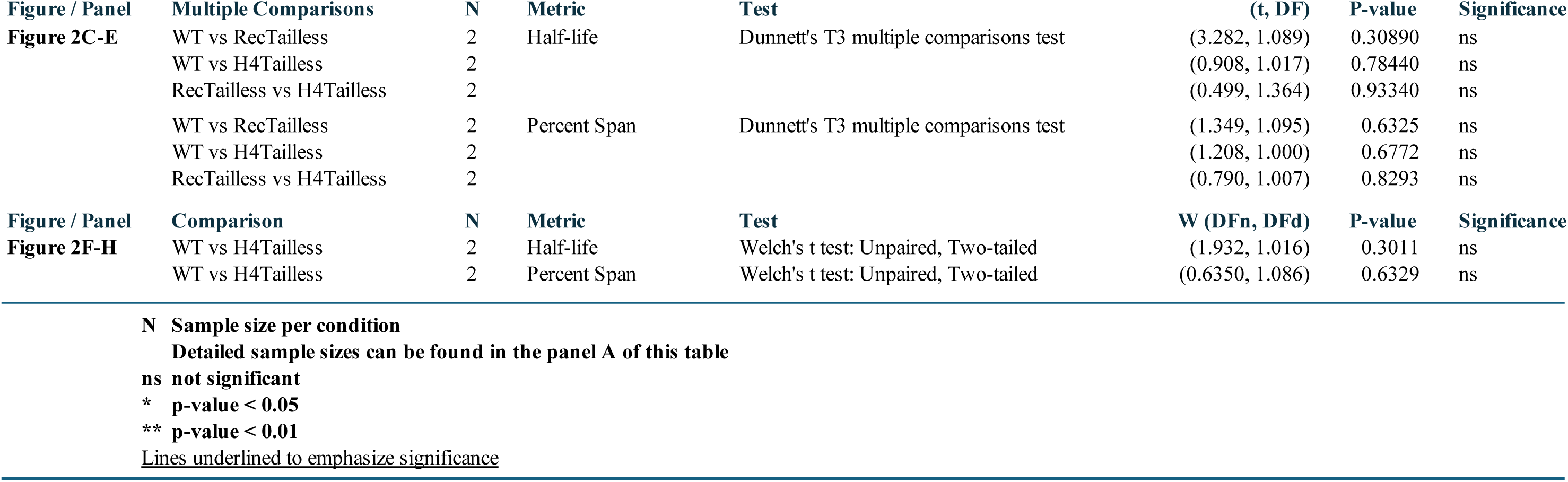
Kinetic parameters, model-selection metrics, and statistical comparisons for SWR1C-mediated H2A.Z exchange on WT, RecTailless, and H4Tailless mononucleosomes under single-turnover and catalytic conditions (Figures 1, 2, and Supplemental Figure 2).

**Supplemental Table 2.**
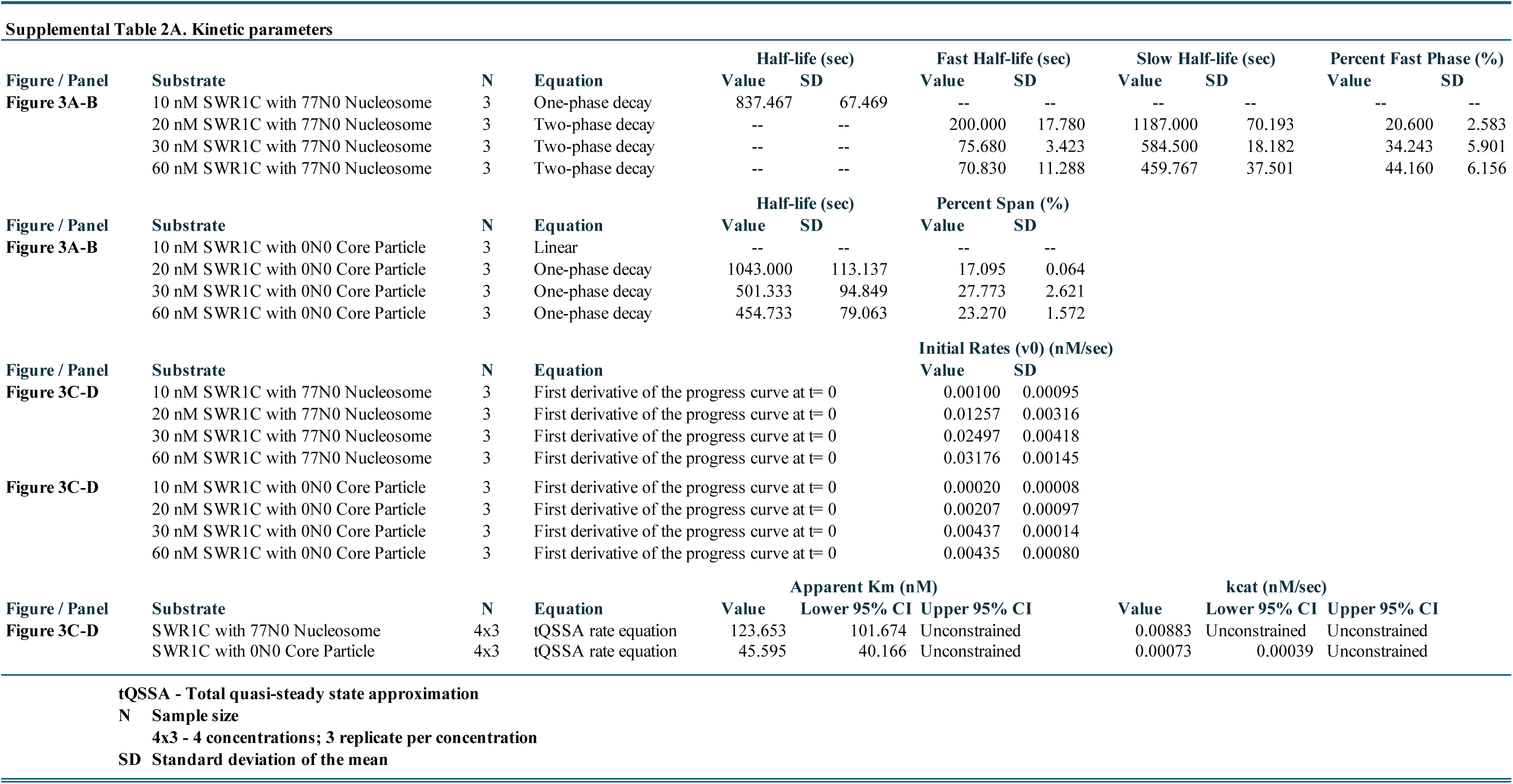
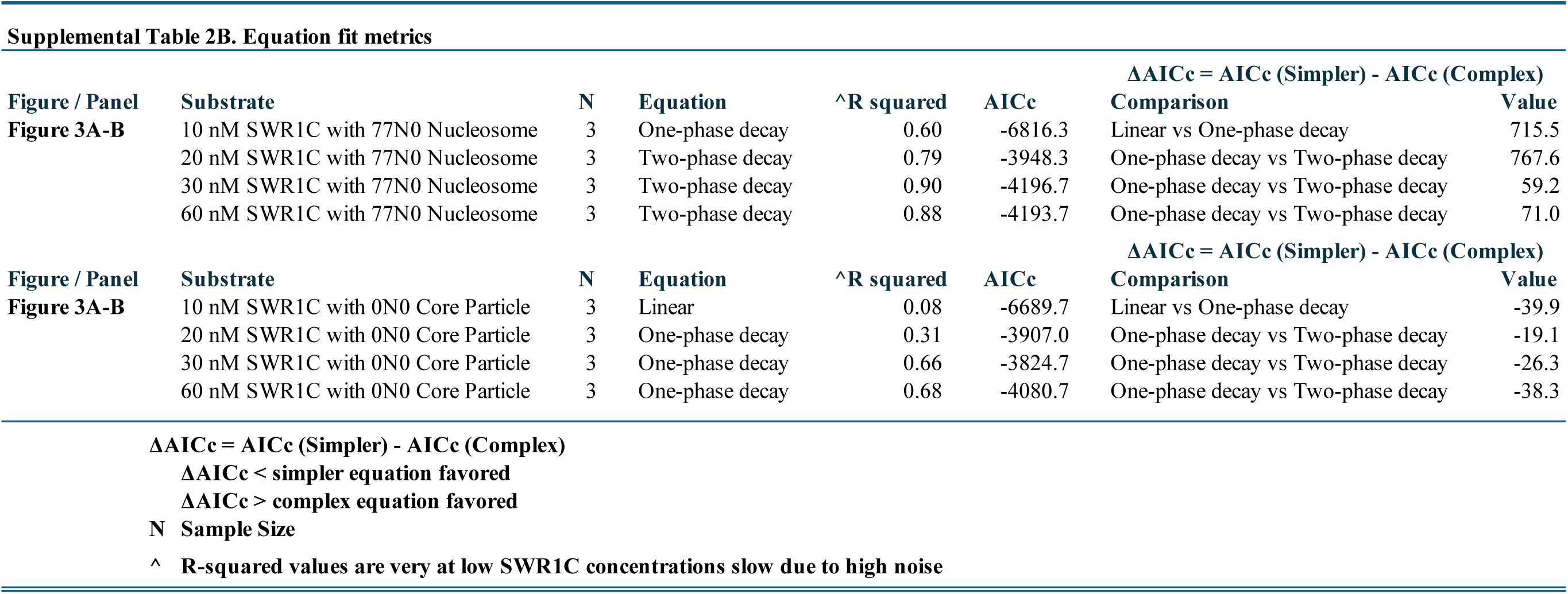
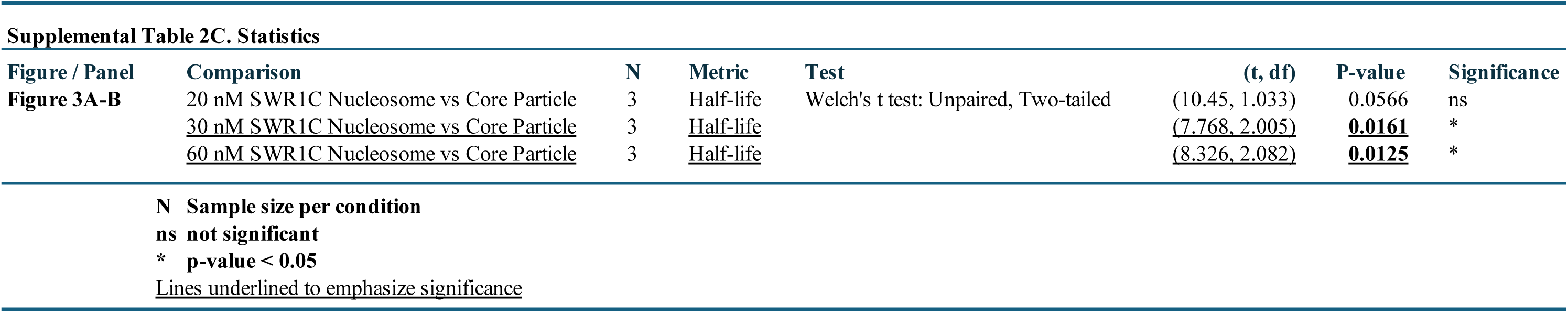
_Linker DNA dependence of SWR1C activity: kinetic fit parameters, progress-curve–derived initial rates, and statistical comparisons for nucleosomes versus core particles across SWR1C concentrations (__Figures 3__)._

**Supplemental Table 3.**
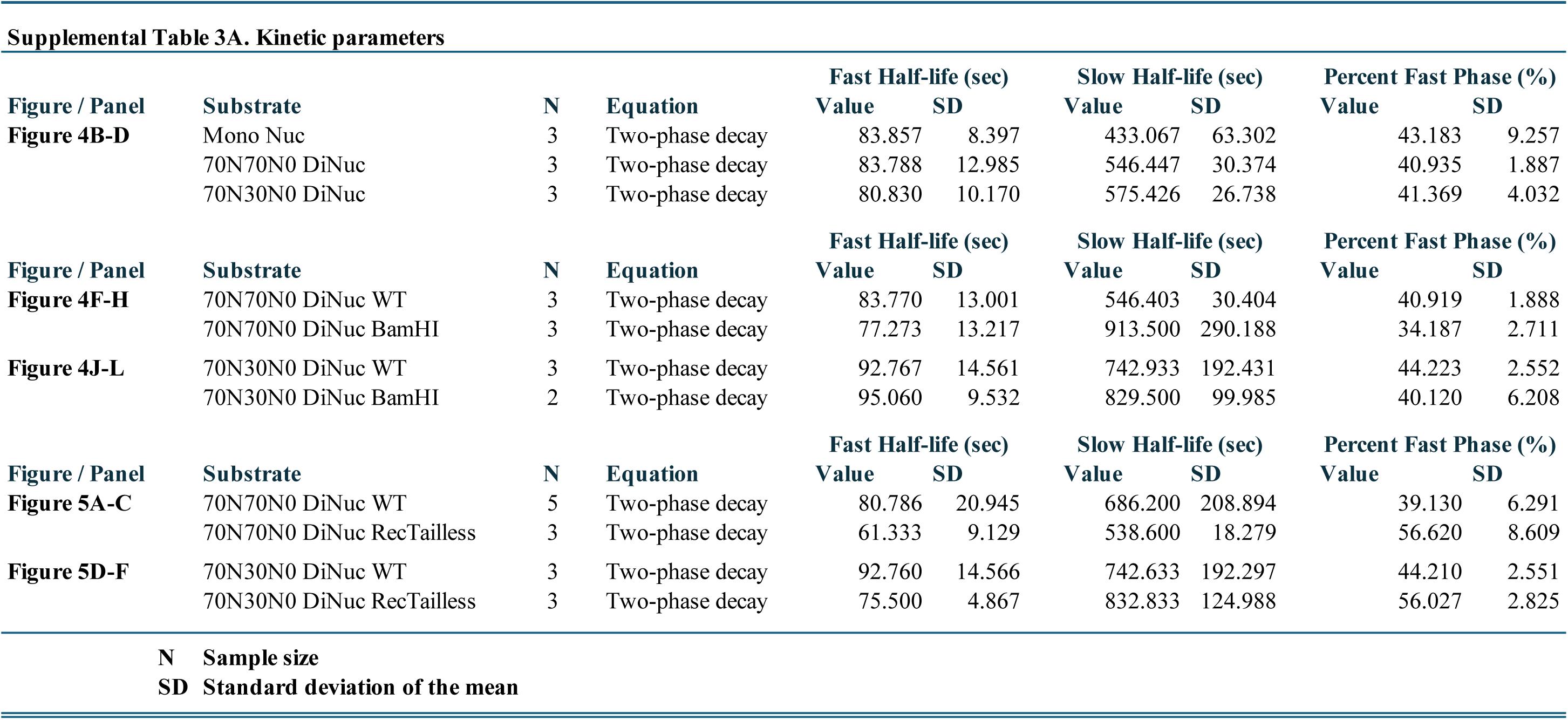
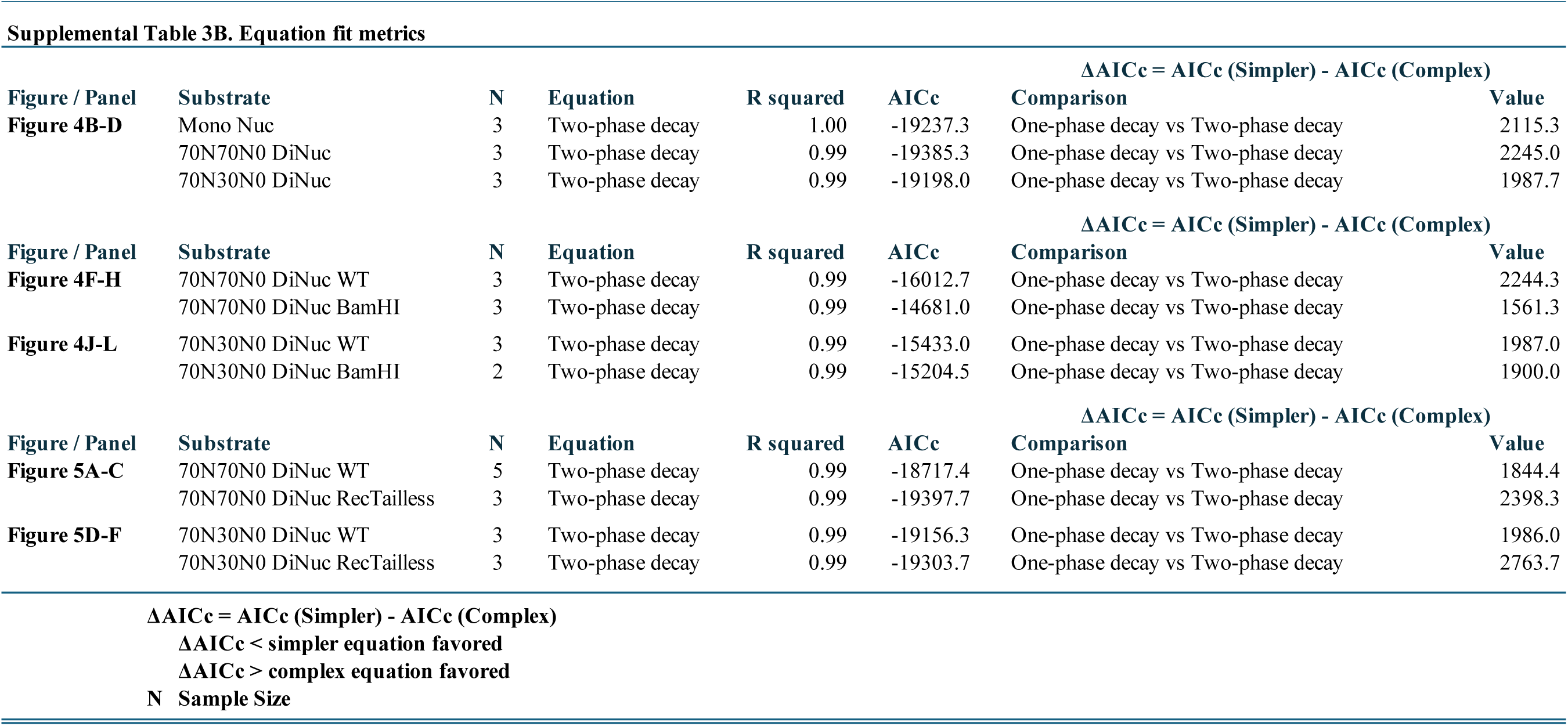
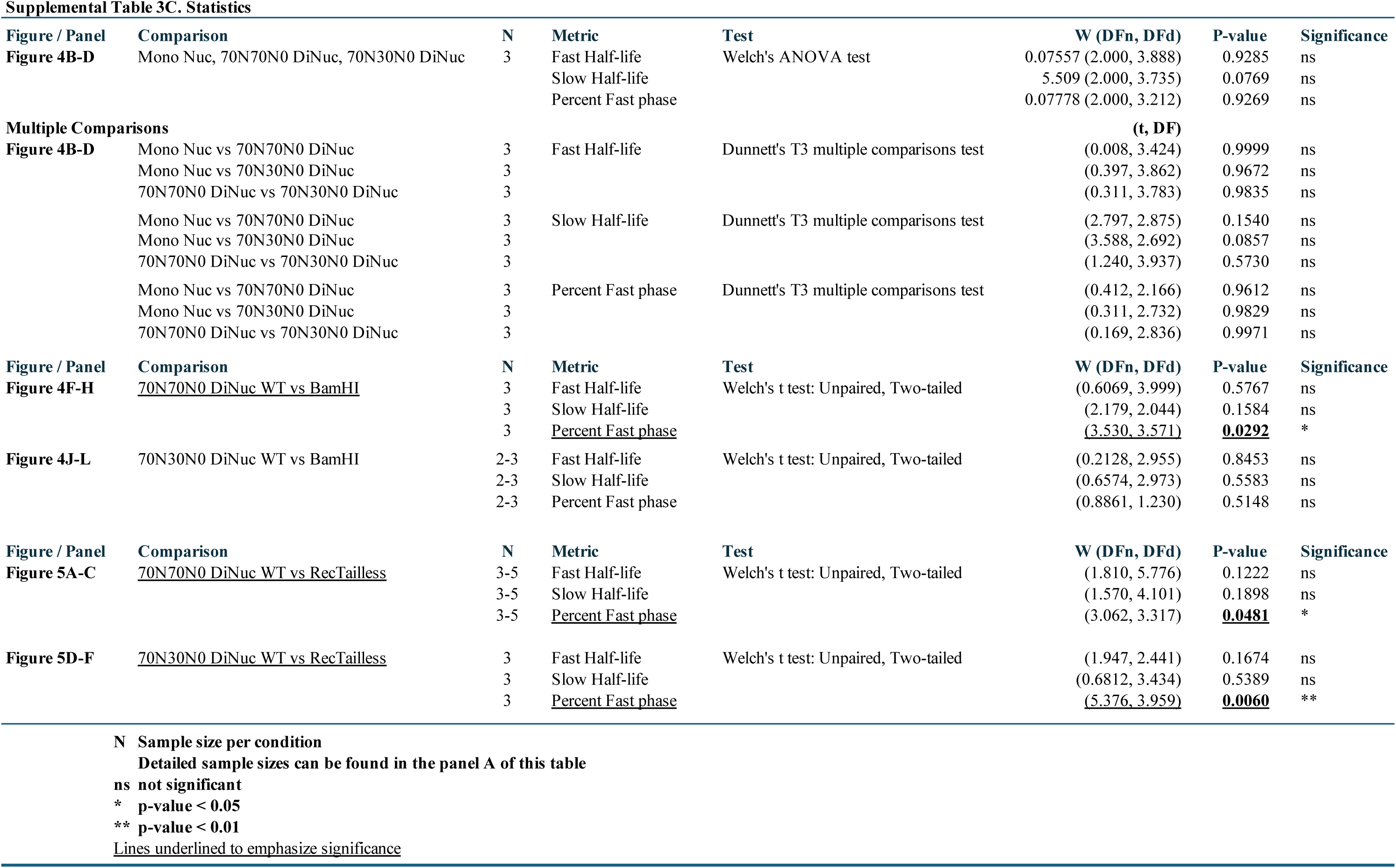
Effects of dinucleosome context and histone tail deletion on SWR1C-mediated exchange: kinetic parameters, model-selection metrics, and statistical comparisons for mononucleosome versus dinucleosome substrates and BamHI-derived mononucleosome products (Figures 4 and 5).

**Supplemental Table 4.**
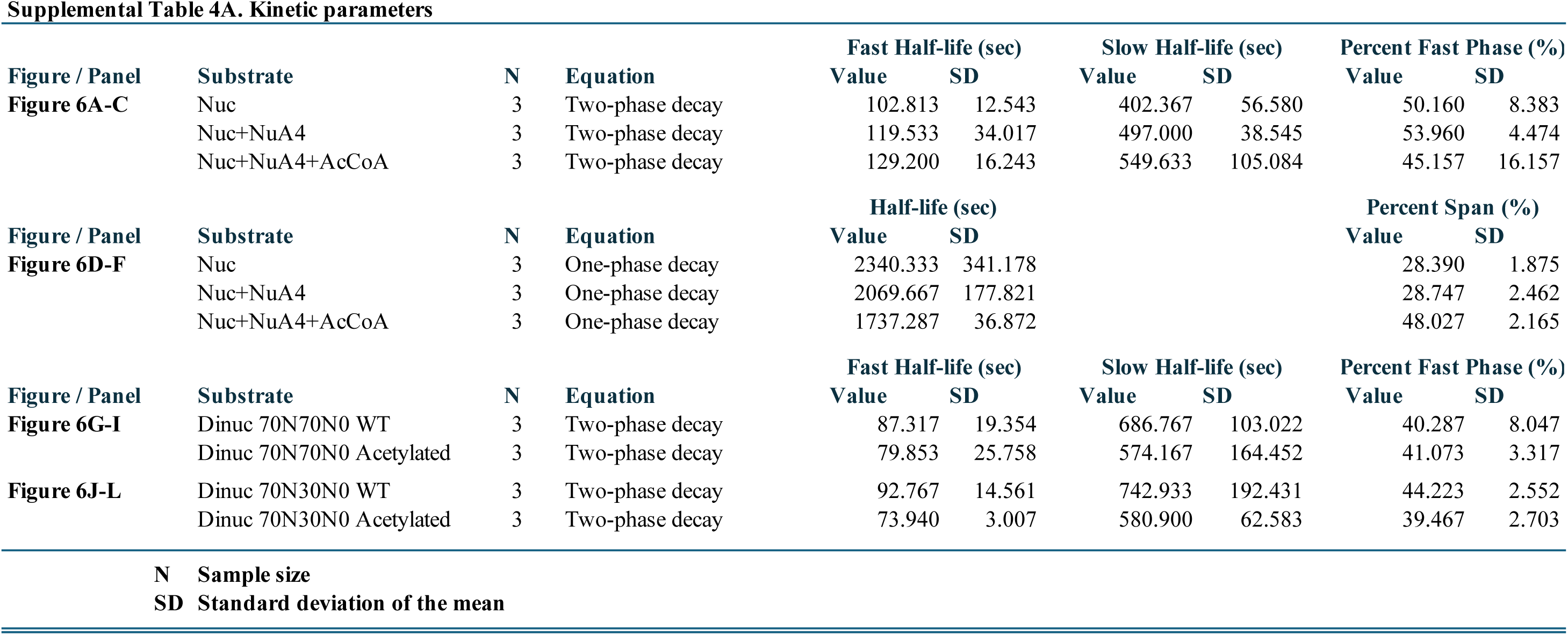
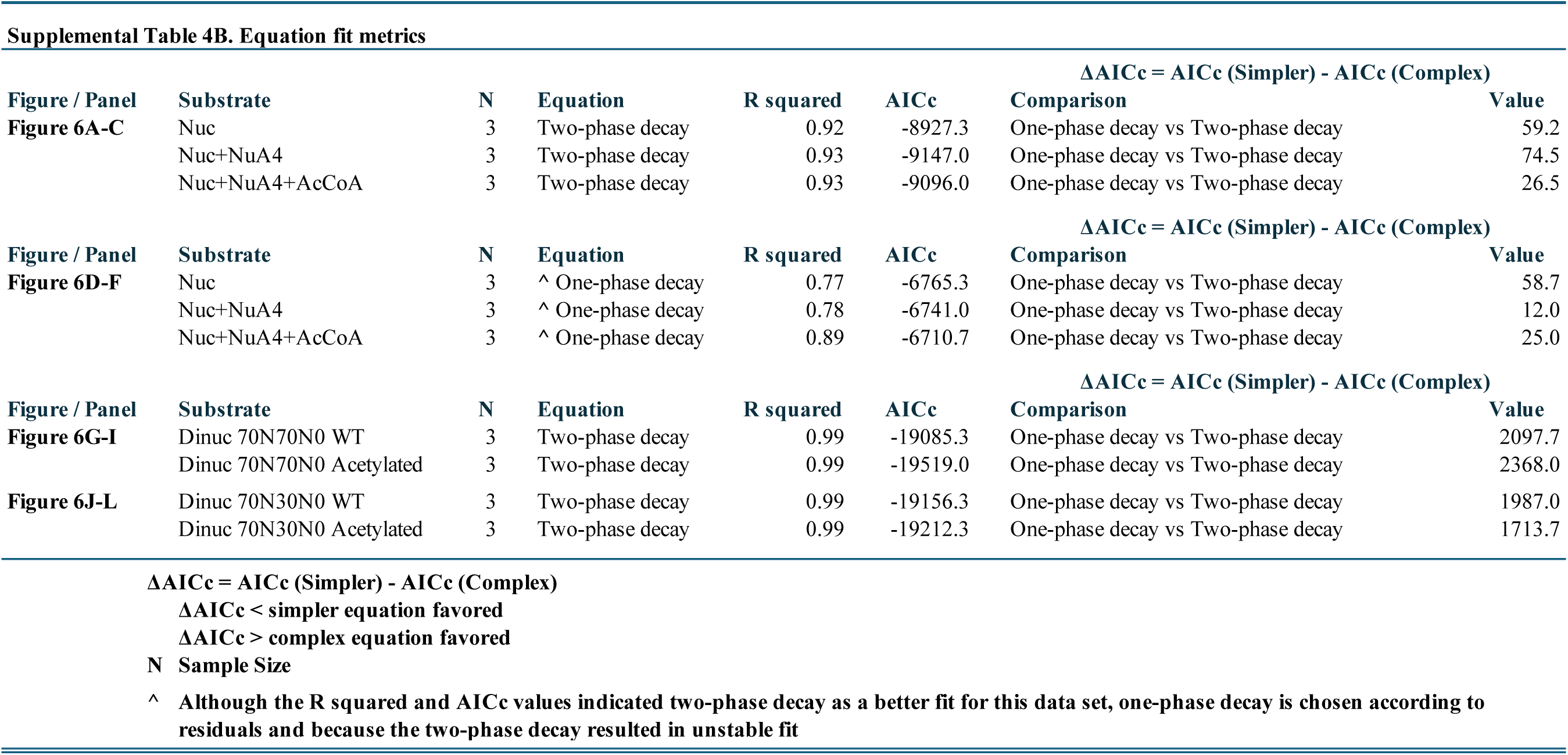
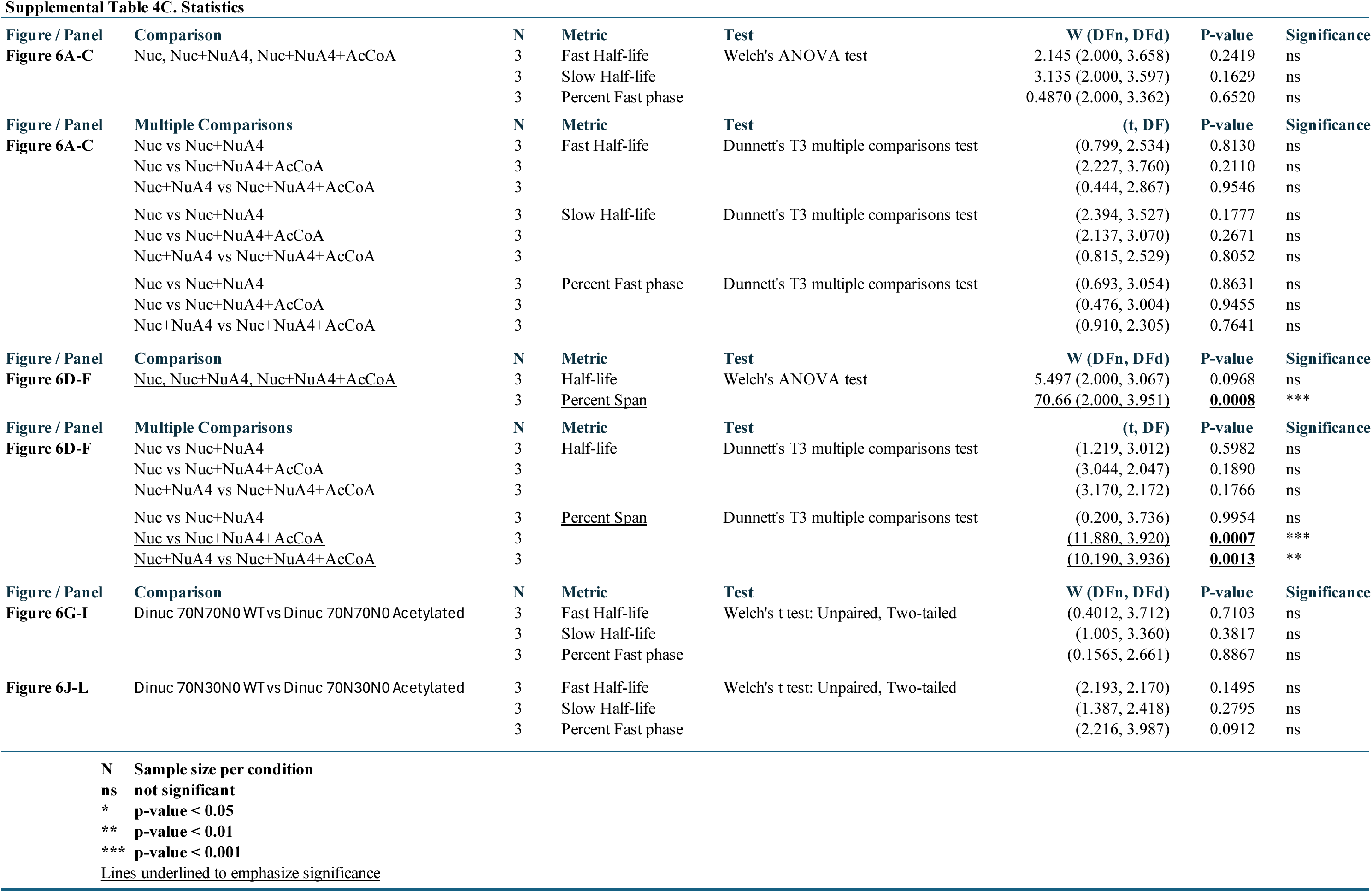
Effects of NuA4-mediated acetylation on SWR1C activity: kinetic parameters, reaction extent metrics, and statistical comparisons for acetylated mononucleosome and dinucleosome substrates, with antibody validation controls (Figure 6).

